# Stochastic transcriptional pulses orchestrate flagellum biosynthesis in *E. coli*

**DOI:** 10.1101/508580

**Authors:** J. Mark Kim, Mayra Garcia-Alcala, Enrique Balleza, Philippe Cluzel

## Abstract

The classic picture of flagellum biosynthesis in *E. coli*, inferred from population measurements, describes a tightly controlled, deterministic transcriptional program. In individual *E. coli* cells, we discover that flagellar promoters are in fact stochastically activated in pulses. Such pulses comprise coordinated ‘on’ and ‘off’ states of promoter activity, each of which can span multiple generations. We demonstrate that this pulsing program obeys the regulatory logic of flagellar assembly, which dictates whether some promoters skip pulses. Remarkably, pulses in this transcriptional network appear to be actually governed by a post-translational circuit. Our results suggest that even topologically simple transcriptional networks can generate unexpectedly rich temporal dynamics and phenotypic heterogeneities.

---

Transcriptional cascades—regulatory motifs where a transcription factor drives the expression of second transcription factor—form the basis for many complex genetic programs that govern cell fate such as division and differentiation (*1-3*). A classic “model” of a transcriptional cascade is the three-tiered gene regulatory network for flagellum synthesis in *E. coli* (**Fig. 1A**) (*4*) which plays crucial roles in locomotion, biofilm formation, surface adhesion, and host invasion (*5-8*). Transcription of the gene *flhDC* (Class I genes) produces the master regulator FlhD_4_C_2_ (henceforth abbreviated as “FlhDC”), which activates the expression of genes involved in the synthesis of the flagellar hook and basal body (Class II genes). One Class II gene encodes the alternate sigma factor FliA, which then activates the expression of genes that encode the flagellar filament and chemotaxis signaling network (Class III genes).

**Figure 1.**
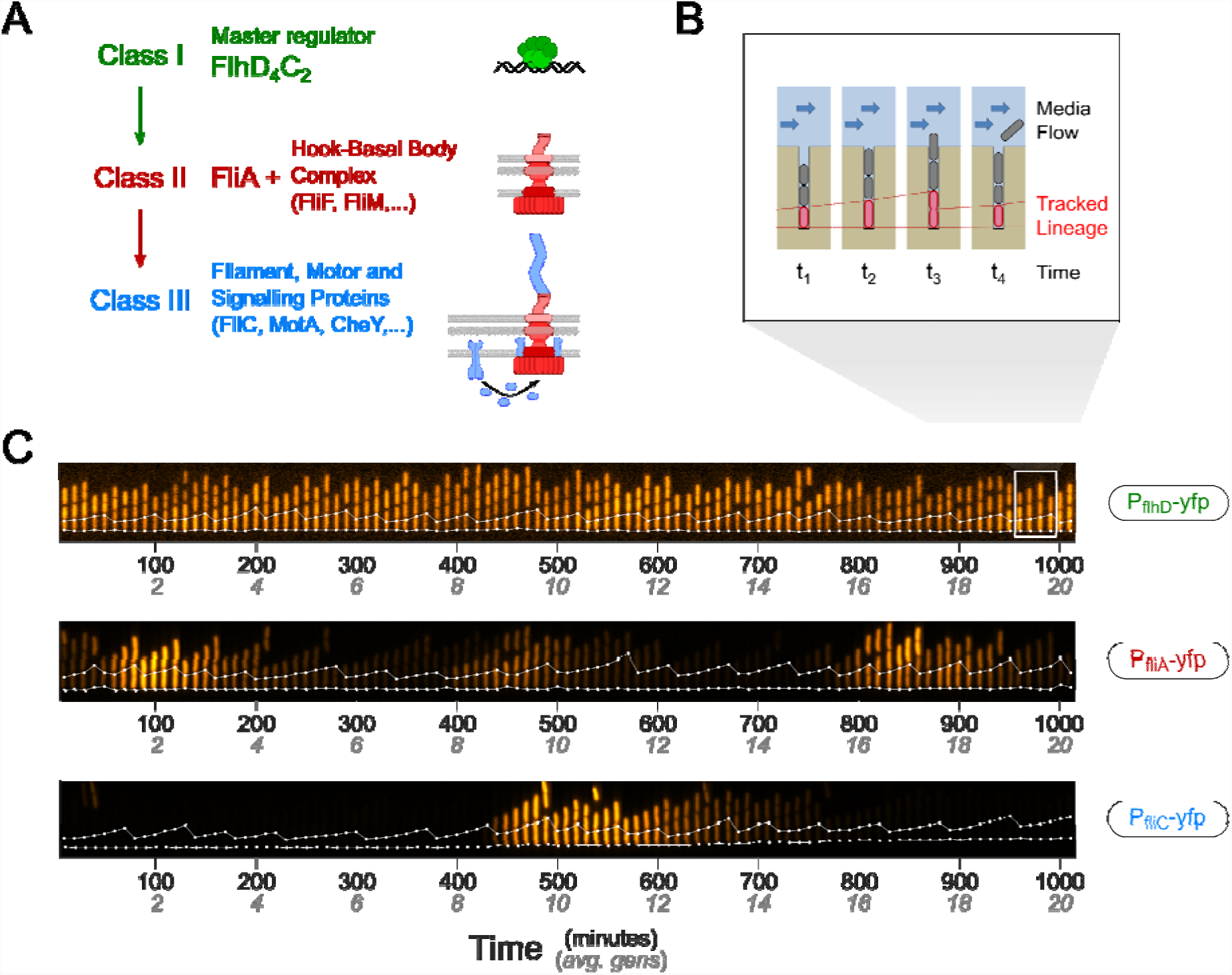
Stochastic pulsing of flagellar promoters: (**A**) The flagellar operons are organized in a transcriptional three-tiered cascade and are accordingly labeled Class I (green), Class II (red) or Class II (blue). For simplicity, our diagram omits various branching and feedback regulatory pathways (such as the anti-sigma factor FlgM). (**B**) A microfluidic device (“mother machine”) constrains cells to grow in narrow linear tracks (**SM D-b**). As the cells grow and divide, they are flushed away at the open end of the tracks by a constant flow of media, which ensures chemostatic growth. We monitor the cell confined at the bottom (the “mother cell”, red) to build a lineage over multiple cell divisions. (**C**) Kymographs illustrating the pulsating dynamics of flagellar promoters. Each kymograph shows fluorescence (falsecolor, orange) from a strain harboring a YFP transcriptional fusion to a specific flagellar promoter from either Class I (top), II (middle) or III (bottom). Cells were grown in MOPS rich defined media (**SM C**) at 34°C and fluorescence images from each cell was acquired every 10 minutes. White dots indicate the location of the mother cell in each frame, which was identified via a constitutively expressed mCherry marker. Cells with the promoter encoding the Class I master regulator (top) are continuously fluorescent. By contrast, Class II (middle) and Class III promoters (bottom) stochastically switch between active and inactive transcriptional states over multiple divisions.

Growing evidence suggests that individual bacteria, even under identical growth conditions, are capable of executing markedly distinct gene expression programs from the rest of the population (*9-14*). However, to date, studies of flagellar gene expression in *E. coli* have largely relied on measurements of gene expression from bulk populations that would mask such single-cell effects (*15-17*). To address this issue, we used a microfluidic device (*18, 19*), which allowed us to monitor flagellar promoter activity in individual cells while maintaining a constant growth environment (**Fig. 1B, Fig. S1, Supplementary Materials, SM section D-d**). In this device, cells are loaded into channels where one end is closed, and the other end is open to microfluidic flow of fresh media. In this way we could dissociate the behavior of the flagellar system from its response to uncontrolled fluctuating environments and analyze with high precision temporal variations taking place in the flagellum regulatory network itself.

Operationally, we constructed *E. coli* strains in which we inserted in the chromosome a copy of a flagellar promoter fused to the coding sequence of a yellow fluorescent protein (YFP) variant mVenus NB (*20*) (**SM B**). For *flhDC*, we also made a cis-insertion of YFP at the end of the *flhDC* transcript to more directly monitor the production of the endogenous *flhDC* transcript. Using the microfluidic device, we could monitor YFP expression in the trapped “mother” cells growing at the bottom of each channel for over twenty generations.

In three distinct experiments, we tracked the activity of promoters from the three different classes (**Fig. 1C**). The promoter of the Class I gene *flhDC* behaved like a standard constitutive promoter with a steady expression over time with small random fluctuations about the mean. However, Class II promoters (exemplified by *fliA*) showed a surprising pulsatile pattern of transcription: cells exhibited long periods of inactivity lasting multiple generations which were suddenly interrupted by several generations of highly promoter activity, before switching back to inactivity. Class III promoters (exemplified by *fliC*) also pulsated, but with inactive periods that were much longer than that observed for Class II promoters. We found that single cells grown in liquid culture also displayed highly heterogeneous flagellar gene expression consistent with the pulsing behavior observed in our microfluidic device (**Fig. S2, Fig. S3**).

To characterize the promoter dynamics more precisely, for each “mother” cell lineage, we computed the promoter activity from the fluorescence signal time series (**Fig. S5; SM D-d**). Although the activity of the Class I promoter fluctuated, the relative variability (as measured by coefficient of variation (CV)) was marginally larger than that of a constitutive promoter (CV_Class1_=1.1 vs CV_Const_=1.06) (**Fig 2A, Fig. S6**). By contrast, Class II and III promoters (**Fig. 2B, C**) showed much larger variations in promoter activity relative to its mean (CV_Class2_=3.6, CV_Class3_=4.4). Remarkably, when we monitored two Class II promoters within the same cell (via CFP and YFP), we found that the pairs pulsated synchronously (**Fig. 2B**). Similarly, we found that two Class III promoters within the same cell also behaved synchronously (**Fig. 2C**). This observation could be confirmed by the high correlation between the CFP and YFP reporters in cells with a pair of Class II or Class III promoters (**Fig. 2B and C bottom; Fig. S7**). Based on this observation, we subsequently focused on the promoters of *flhDC, fliF,* and *fliC* as representative promoters of Class I, II and III respectively.

**Figure 2.**
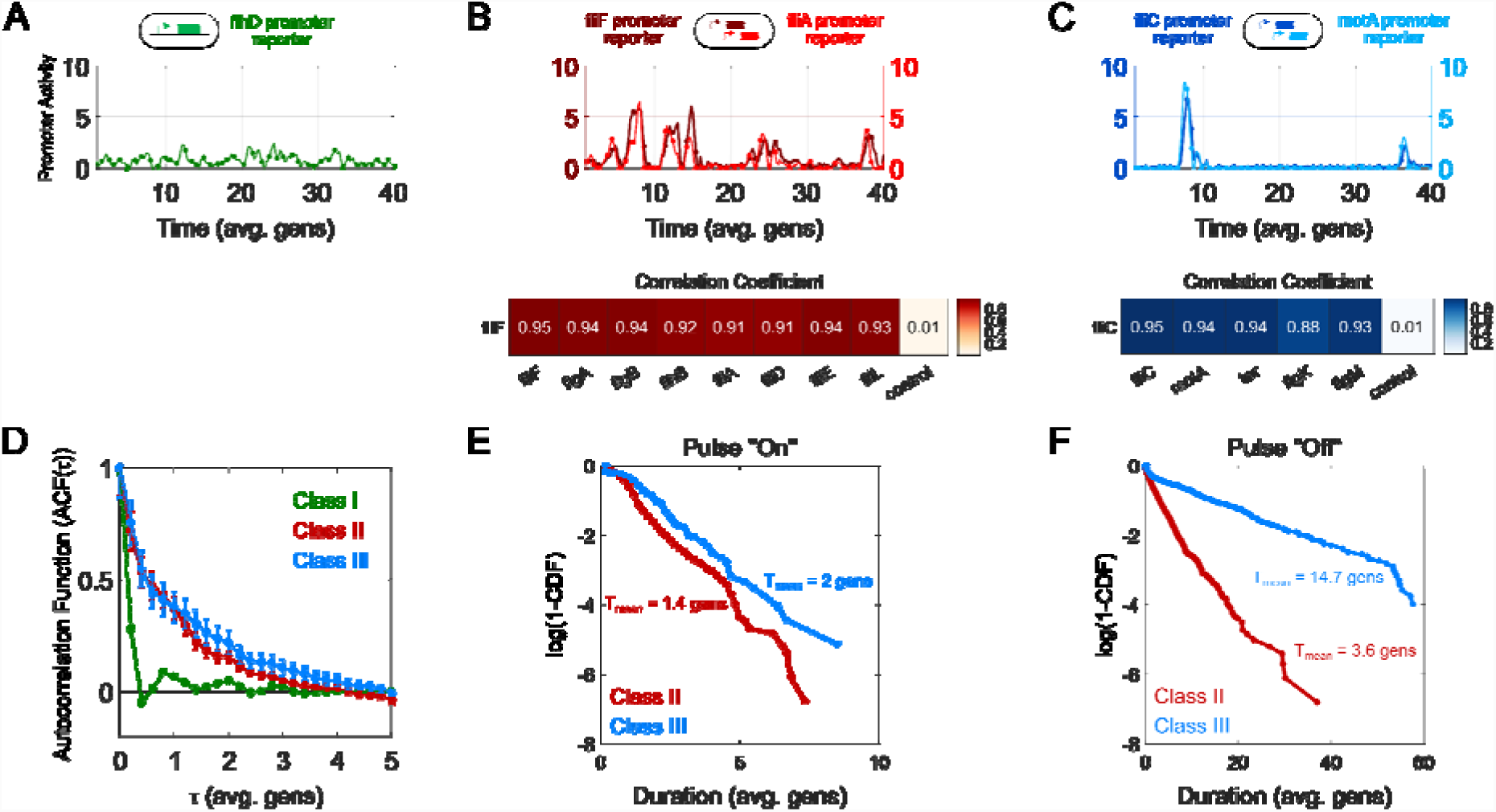
Promoters across classes show distinct dynamics but pulse simultaneously within their own class (II or III). (**A**) Typical activity of the sole Class I promoter, flhD, which controls the expression of the master regulator (green). Promoter activity is quantified by taking the cell-growth corrected time-derivative of the associated fluorescence signal (**SM D-d**). (**B**) (Top) Activity of fliF (dark red) and fliA (bright red) promoters within the same cell, representative of Class II pulsing dynamics. (Bottom) Correlation between two Class II gene reporters in the same cell as determined by flow cytometry (**SM D-a**). Each strain harbors a reference reporter consisting of the fliF promoter and CFP and a second Class II promoter fused to YFP. The control promoter is a synthetic constitutive promoter. (**C**) Activity of fliC (dark blue) and motA (light blue) promoters within the same cell, representative of Class III pulsing dynamics. (Bottom) Correlation between two Class III gene reporters in the same cell as determined by flow cytometry. Similar to (B), each strain harbors a reference reporter consisting of the fliC promoter and CFP and a second Class III promoter fused to YFP. The control promoter is again a synthetic constitutive promoter. (**D**) Normalized autocorrelation function of flagellar promoter activity of Class (green), II (red), and III (blue), estimated from the activity of flhD, fliA, and tar promoters, respectively (**SM D-d**). For each promoter, the autocorrelation was estimated from 50 lineages, each at least 40 generations long (**E**) Cumulative distribution of the pulse “on” durations, plotted as log(1-CDF). Shown are the distributions for fliF (Class II, red) and fliC (Class III, blue). Durations were estimated from 50 lineages, each at least 60 generations long. (**F**) Cumulative distribution of the pulse “off” durations, plotted as log(1-CDF). Shown are the distributions for fliF (Class II, red) and fliC (Class III, blue) Durations were estimated from 50 lineages, each at least 60 generations long.

The fast stochastic fluctuations in *flhDC* transcription were dynamically distinct from the pulses in Class II and Class III promoters as indicated by the autocorrelation functions of these three promoters. While the autocorrelation function of *flhDC* promoter activity decayed rapidly, the Class II and Class III pulses showed longer correlation times, consistent with the longer timescale of the pulses (**Fig. 2D**). To quantify the duration of the pulses, we defined an operational threshold based on the detection limit of our reporters and categorized the promoter as being “on” or “off” based on whether the activity was above or below that threshold. The “on” duration of the pulses were approximately exponentially distributed for both Class II and Class III pulses with similar mean durations spanning 1-2 cell generations (**Fig. 2E**). The “off” durations were also approximately exponentially distributed—however, the average “off” durations of Class II promoters were significantly shorter (~4 generations) compared to Class III promoters (~14 generations) (**Fig. 2F**). We note that these values may slightly underestimate the true “on” and “off” durations since we often observed long clusters of pulses interrupted by short periods of inactivity. Overall, our results suggest that despite being coupled in a linear cascade, each class of flagellar promoters has a distinct dynamical behavior.

How might transcription of *flhDC* give rise to pulsatile behavior of Class II promoters? Given that Class II (and III) promoters appear to pulse in spite of relatively steady *flhDC* transcription, we first decided to examine whether any endogenous transcriptional or translational regulation of *flhDC* was necessary for Class II pulses. Various studies have established that transcription of *flhDC* is regulated by multiple pleiotropic transcription factors (*21-24*). Additionally, small RNAs (sRNAs) have been shown to regulate the translation and/or lifetime of the *flhDC* mRNA by binding to the 5’ untranslated region (5’-UTR) (**Fig. 3A, left**) (*25*). To bypass this regulatory complexity, we replaced the ~2kb region directly upstream of the endogenous *flhDC* coding sequence with synthetic sequences encoding constitutive promoters of various strength (“the Pro series”) (*26*). We also modified the DNA sequence corresponding to the 5’ UTR of the *flhDC* mRNA with a synthetic ribosomal binding site (RBS) sequence derived from the T7 phage. Consequently, in these strains, expression of FlhDC was isolated from any known transcriptional and translational regulators (**Fig. 3A, right**). Surprisingly, when expression of *flhDC* was coupled to synthetic promoter Pro4, downstream Class II promoters began pulsing similar dynamics and amplitudes than those observed in the presence of the native promoters (**Fig. 3B, C**). Thus, Class II pulses do not require endogenous transcriptional or transcriptional regulation of the master regulator.

**Figure 3.**
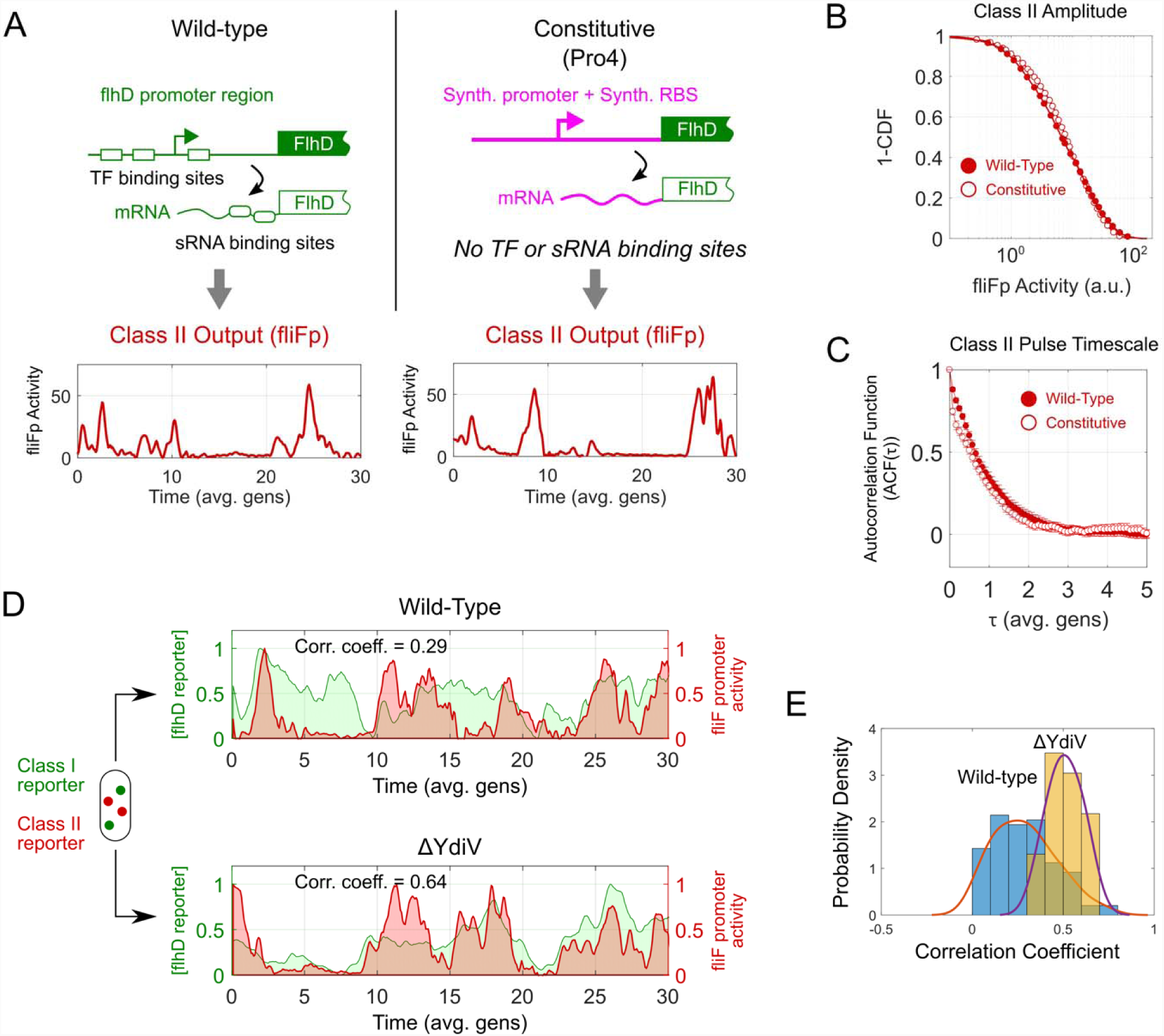
Class II pulses do not require transcriptional or translational endogenous regulation of the master regulator. (**A**) **Top,** Endogenous expression of the master regulator FlhDC is driven by the class promoter, whose regulation is controlled by multiple transcription factors (TF)(41). In addition, translation of FlhDC is regulated by several small RNAs (sRNAs) interacting with the 5’ untranslated region (UTR) of the FlhDC transcript (**top left**). We replaced the native Class I promoter with a synthetic constitutive promoter (Pro4)—additionally, we altered the 5’ UTR so that the synthetic ribosoma binding site (RBS), which drives FlhDC translation is insensitive to sRNA regulation (**top right**). **Bottom,** typical Class II promoter dynamics (fliF promoter, red), wild-type (**bottom left**) and “constitutive”(**bottom right**) FlhDC promoters. (**B**) Cumulative distribution function (plotted as 1-CDF) of Class I promoter activity amplitudes from wild-type (solid red circle) and “constitutive” strains (open red circle) For each strain, we analyzed 50 lineages, each at least 30 generations long. (**C**) Normalized autocorrelation function of Class II promoter activity, wild-type (dark red), synthetic “constitutive” (light red). For each strain, we analyzed 50 lineages, each at least 30 generations long. (**D**) Simultaneous measurements of fluorescence signal from the Class I reporter (green) and activity of the Class I promoter fliFp (red) within the same cell. The fluorescence signal from the Class I reporter is a proxy for the concentration of proteins produced from the FlhDC promoter while the Class II promoter activity is the time derivative of the fluorescence signal from the Class II reporter which is a proxy for transcription from that promoter. Typical examples of wild-type (upper) and ΔYdiV cells (lower) along with the Pearson correlation coefficient for the Class I and Class II signals (**SM D-d**). For ease of visualization, each signal is normalized so that the minimum value of the signal is 0 and maximum value of the signal is 1. Pearson correlation coefficient was computed on the raw data prior to normalization. (**E**) Normalized histogram showing distribution of correlation coefficients from 100 lineages for wild-type (blue) and for ΔYdiV (yellow) strains. Solid lines, kernel density approximations of those distributions (red and purple respectively). Each lineage is at least 30 generations long.

We then asked whether post-translational regulators of FlhDC activity could influence the dynamics of the observed class II pulses. To this end, we turned to the anti-FlhDC factor YdiV ((*27, 28*). Although the regulatory role of YdiV was well characterized in *Salmonella* (*28*), both the expression and regulatory significance of the *E. coli* homolog had previously been questioned (*27, 29*) (**SM F**). To test whether YdiV plays a role in Class II pulses, we first generated a strain harboring a cis-insertion of YFP at the 3’ end of the *flhDC* transcript and a trans-copy of the *fliF* promoter fused to CFP in another chromosomal locus. In wild-type cells, we observed that the fluctuations in YFP fluorescence levels, a proxy for the FlhDC concentration, occasionally coincided with Class II activity but that this correlation was very weak (R=~0.3) (**Fig. 3D, top; Fig. 3E**). Upon deletion of YdiV mutant strains, we found that Class II promoter activity became considerably more correlated (R~0.6) with the YFP fluorescence (**Fig. 3D, bottom; 3E**). Taken together, our results suggest that random transcriptional “noise”—rather than active transcriptional or translational regulation—of *flhDC*, coupled with post-translational regulation of FlhDC by YdiV, generate the stochastic pulses of Class II promoter activation. Interestingly, we found that flagellar genes such as FliZ and FliT which have been implicated in post-translational regulation of FlhDC in *Salmonella*, appeared to have only a minor effect on flagellar pulses in *E. coli* (**SM F; Fig. S9**).

We also examined how pulses in Class II promoters might be transmitted to Class III promoters. Using a similar approach we used to study the transcriptional cascade from Class I to Class II, we generated a strain harboring both a copy of the Class II (*fliF*) promoter driving CFP and a copy of the Class III (*fliC*) promoter driving YFP. We then compared the Class II fluorescence signal, a proxy for the concentration of proteins produced from Class II promoters, to the activity of a Class III promoter. We found that pulses in Class II genes were not always accompanied by pulses in Class III genes, which skipped in a stochastic manner some of the Class II gene pulses (**Fig. 4A**). However, all Class III pulses were always accompanied by Class II pulses (**Fig. 4A**).

**Figure 4.**
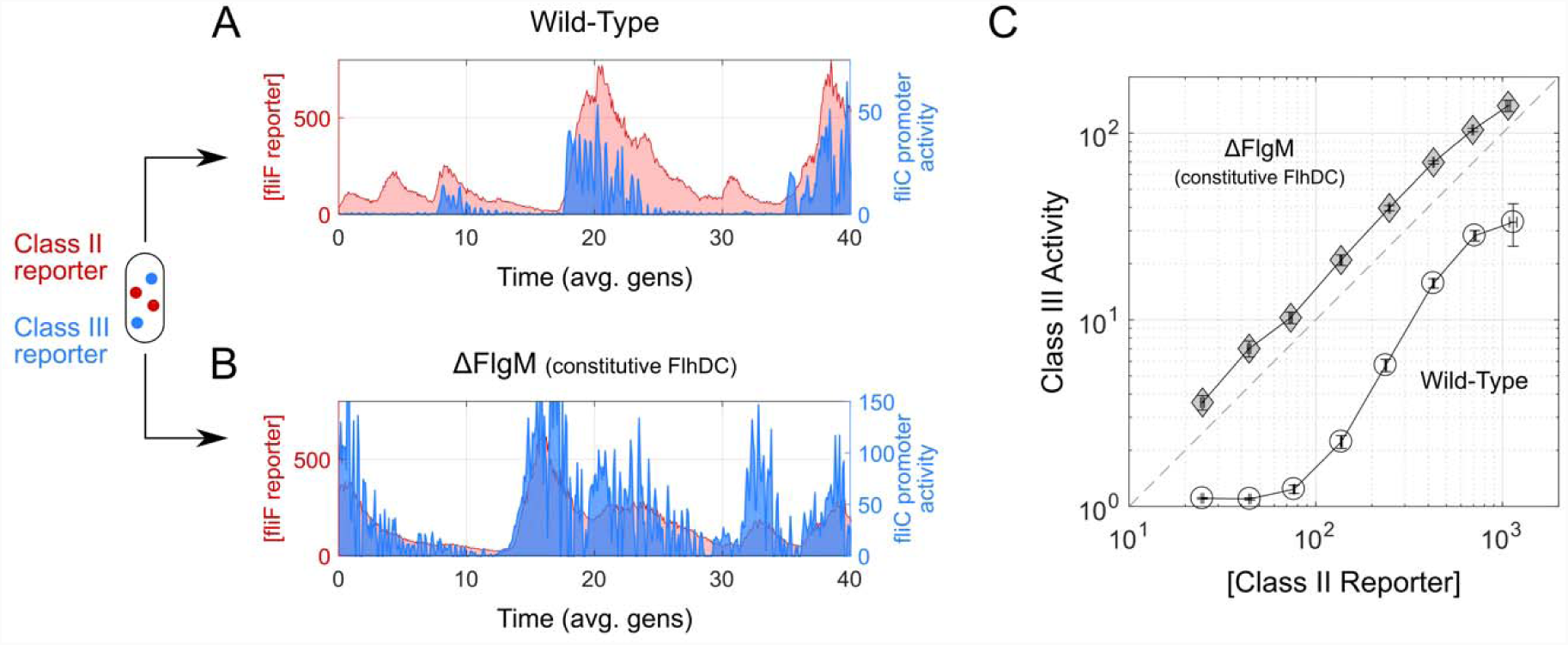
Modulation of Class III pulses by post-translational regulation of the alternative sigma factor, FliA. (**A and B**) Paired measurements of Class II and Class III within the same cells. We compared fluctuations in Class II fluorescence signal (red, fliFp), a proxy for the concentration of proteins produced from Class II promoters, to the activity of a Class III promoter (fliCp, blue). Typical examples, (**A**) wildtype cells, and (**B**) a ΔFlgM mutant that constitutively expresses the master regulator, FlhDC, to bypass any transcriptional feedback (**SM A**). (**C**) Mean Class III activity as a function of the Class II reporter concentration in cells harboring fliF and fliC promoter reporters. Binned average of single cel measurements, wild-type (circles) and ΔFlgM (diamonds) strains; ΔFlgM mutant, same as in (B). For each strain, we analyzed 25 lineages, each at least 50 generations long.

A well-characterized checkpoint regulates the transcriptional cascade from Class II to Class III promoters: Transcription of Class III genes is regulated by the Class II gene, the alternative sigma factor FliA (*30, 31*). However, another Class II gene, FlgM, binds to FliA and acts as an inhibitor (*32*). Thus FliA remains initially inactive. When the basal body of the flagellar motor (encoded by the other class II genes) is assembled, it acts as a secretion system and exports FlgM from the cytoplasm (*33, 34*). This export results in the activation of FliA. We hypothesized that this checkpoint pathway, and more directly FlgM, might be responsible for the “skipping” behavior of Class III pulses.

To test this hypothesis, we simultaneously monitored the pulsating dynamics of the Class II (fliFp) and Class III (fliCp) promoters in a FlgM knockout strain (ΔflgM). In this mutant strain, Class III promoters pulsed deterministically whenever Class II genes pulsed (**Fig. 4B**). To better quantify this relationship, we plotted the mean activity of the Class III promoter as a function of the Class II reporter concentration. In wild-type cells, we observed a sigmoidal relationship, which is consistent with the idea that a critical concentration of Class II gene products is necessary for cooperative assembly of the basal body and export of FlgM, which in turn frees the sigma factor FliA for activation of Class III genes. By contrast, in the ΔflgM mutant, the Class II-III relationship became linear, suggesting that Class III pulses now simply mirror Class II pulses (**Fig. 4C**).

Finally, we decided to examine how the dynamics of the flagellar network might change if we varied the mean levels of *flhDC* transcription. Using the previously described synthetic Pro-series promoters, we measured the Class II (*fliF*) promoter activity as a function of a various levels of Class I (*flhDC*) expression (*26*). Given our observation that the inherent noise of these synthetic promoters can lead to large changes in Class II promoter activity (**Fig. 3**), rather than averaging all the single cell data for a given strain, we divided Class I reporter levels into five logarithmically-spaced bins. Then, for each bin we plotted the mean input (i.e. Class I) against the mean of the corresponding output (i.e. Class II) (**SM D-d**). This procedure gave us greater resolution into how the Class II promoter activity changes as a function of relatively small changes in Class I. The resulting plot is analogous to a classic “doseresponse” relationship.

We first examined the “dose-response” relationship in the ΔYdiV mutant to eliminate potentially confounding effects of the anti-FlhDC factor YdiV. Unexpectedly, we discovered that in the resulting dose-response relationship, the transition from inactive to active Class II occurs at a higher levels of Class I than the transition from active to inactive Class II suggesting the existence of a hysteretic cycle (**Fig. 5A**). Additionally, this result contrasts the circuit in *E. coli* to that in *Salmonella* where YdiV plays an essential role in generating bistability and hysteresis (*35*) (**SM F**). In *E. coli*, the presence of YdiV in fact appears to “scramble” the hysteresis (**Fig. 5B**). At levels of *flhDC* production previously corresponding to the “bistable” region (e.g. Pro4), the restoration of YdiV appears to erode this bistability and cause Class II activation to become more smeared (i.e. more heterogeneous) (**Fig. 5B**).

**Figure 5.**
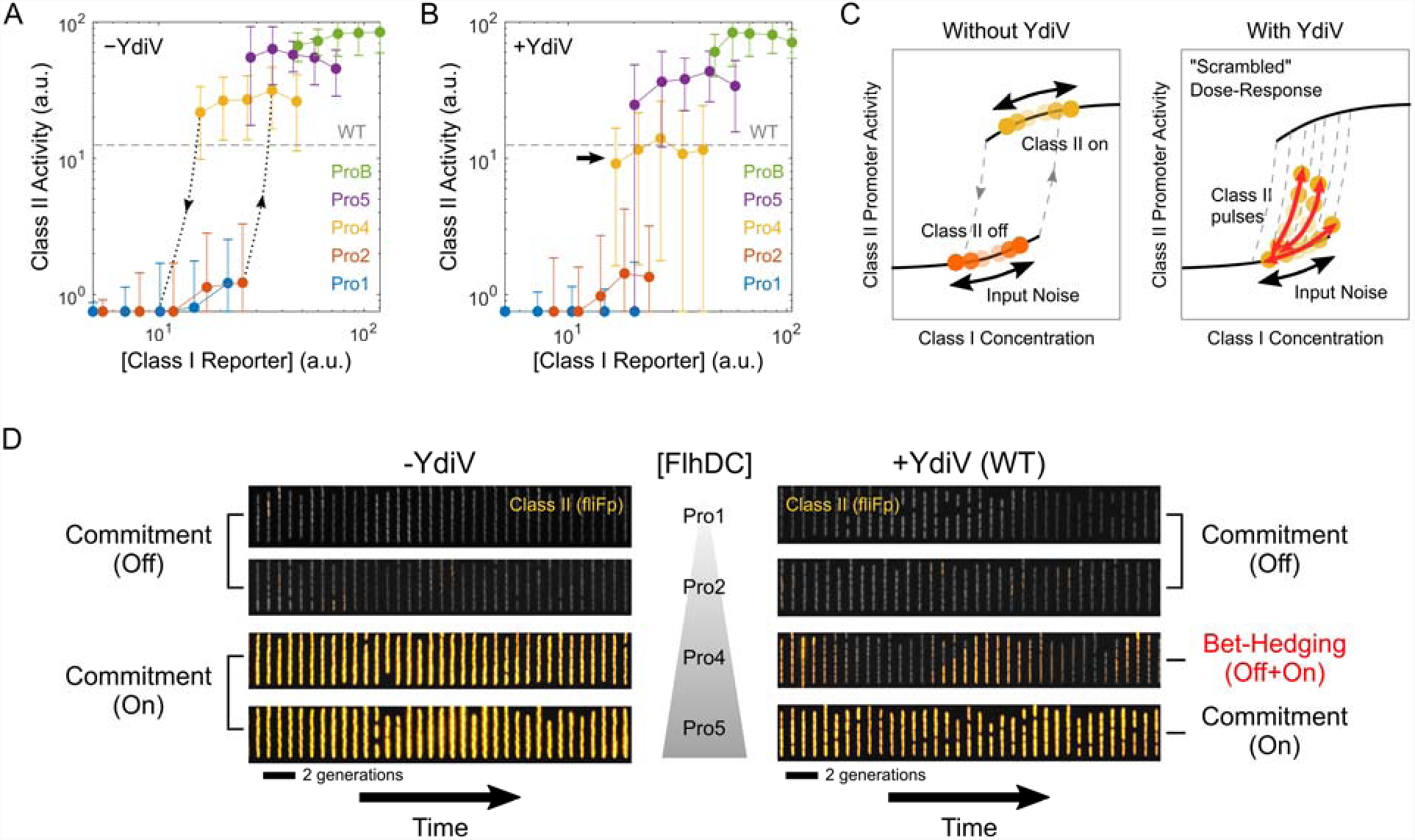
Model of flagellar regulation using a hysteretic switch. (**A and B**) Input-output relationshi pbetween Class I and Class II. We plotted Class II (fliFp) promoter activity (**SM D-d**) as a function of a wide range of Class I levels using synthetic promoters (Pro1-5, ProB). For each strain associated with a different synthetic promoter, we divided Class I reporter levels into 5 logarithmically-spaced bins. From each bin we plotted the mean input (i.e. Class I) against the mean of the corresponding output (i.e. Class II). For ease of visualization, all Class II promoter activities below mean cellular autofluorescence were set to be equal to the autofluorescence value. (**A**) ΔYdiV mutant strains, and (**B**) strains with wildtype YdiV levels. Error bars indicate standard deviation of Class II activity. In (**A**), dashed lines with arrows are guides for the eyes to delimit the hysteretic behavior in ΔYdiV mutants where Class II activity appears to switch abruptly. In (**B**), black arrow highlights the strain (Pro4) whose Class II activity shows the greatest difference with that of ΔYdiV mutants. For each strain in (**A**) and (**B**), we analyzed at least 25 lineages, each at least 20 generations long. (**C**) Schematic illustration of the Class I-Class II dose-response relationship with (right) and without (left) YdiV. (Left) In the absence of YdiV, hysteresis causes Class I to remain “on” (yellow) or “off” (“orange”) in spite Class I transcriptional noise (“input” noise). (Right) YdiV causes the hysteresis to be “scrambled” causing some transcriptional fluctuations to result in large changes in Class II activity (“pulses”). (**D**) Commitment versus bet-hedging behavior in ΔYdiV and wildtype cells. Each strip is a typical kymograph of a strain harboring a Class II promoter reporter expressing FlhDC from synthetic promoters Pro1, Pro2, Pro4 or Pro5. For visual aid, a constitutive marker (gray) was merged with the Class II reporter fluorescence (orange/yellow).

In effect, our observations suggest that in the absence of YdiV, the hysteresis tends to prevent small variations in FlhDC production from drastically altering the state of Class II activation (**Fig. 5C, left**). In effect, cells tend to be committed to keeping Class II “on” or “off”. If YdiV stochastically weakens the hysteresis, this would allow noise in FlhDC production to occasionally trigger an ultrasensitive transition between the two states—i.e. cells could periodically “escape” the hysteresis and become active or inactive (**Fig. 5C, right**). Such a behavior would manifest itself as pulses of in Class II activation overtime. Overall, our results suggest that wild-type circuit appears to be particularly optimized, by the action of YdiV, to generate pulses.

What might be the functional consequence of such a circuit? In the absence of YdiV, increasing FlhDC expression would cause *E. coli* cells previously not expressing any flagellar genes to abruptly commit to flagellar synthesis—such a “hypersensitive” behavior would be beneficial for responding to small changes in the environment but would also force cells to commit to a genetic program for a very long time (**Fig. 5D, left**). By contrast, with YdiV, wild-type *E. coli* can use an “compromise” solution where it can sample between active and inactive Class II expression if expression of FlhDC is intermediate (**Fig. 5D, right**).

Taken together, our results suggest that although the core architecture of the flagellar network appears to be a straightforward linear transcriptional cascade, it exhibits a surprisingly rich and complex dynamical behavior—most notably a “pulsatile-regime” where continuous *flhDC* expression yields spontaneous switching between active and inactive states of flagellar transcription. Unlike transcriptional bursts, which are typically shorter in time scale (<1 generation), flagellar gene pulses span longer time periods to allow for full activation of the cascade (as evidenced by Class III activation), which suggests that pulses might indeed be functional rather than simple “noise”.

The flagellum plays an essential role in chemotaxis and the initiation of biofilms (*36, 37*). However, its synthesis is associated with a sizeable growth cost (*38*). Since many conditions which trigger *E. coli* flagellar biosynthesis are also associated with decreased nutrient availability, it is then plausible that in order to minimize cost, bacteria have developed such pulsating dynamics as a bet-hedging strategy. This strategy would yield a mixed population with cells favoring either growth or flagellum synthesis. Similarly, pulses in the flagellar system might allow bacteria seeking to colonize host-organisms to circumvent or minimize immune responses (*8, 39, 40*). In general, our results lead us to speculate that under certain environmental conditions *E. coli* can operate the flagellar synthesis network in a “pulsatile” mode of operation, which allows cells to sample different phenotypes over time.

## Acknowledgements

We thank M. Cabeen and J. Paulsson for sharing an early version of the microfluidic mother machine with our groups, M. Cabeen, N. Lord, T. Norman and S. Canas Duarte for technical help with the microfluidic device, R. Losick and his lab for their microscope, and N. Kleckner, J. Paulsson and K. Gibbs for their helpful discussions and feedback. This work was performed in part at the Center for Nanoscale Systems (CNS), a member of the National Nanotechnology Infrastructure Network (NNIN), which is supported by the National Science Foundation under NSF award no. ECS-0335765. CNS is part of Harvard University. This work was supported by grant 1615487 from the NSF.

## Author Contributions

JMK and PC conceived the study. JMK designed and generated genetic constructs, performed experiments, and analyzed data with help from MG and EB. PC supervised the project. JMK and PC wrote the manuscript.

## References

1. A. Martinez-Antonio, S. C. Janga, D. Thieffry, Functional organisation of Escherichia coli transcriptional regulatory network. J Mol Biol 381, 238–247 (2008).

2. J. Bahler, A transcriptional pathway for cell separation in fission yeast. Cell Cycle 4, 39–41 (2005).

3. D. W. Allan, S. Thor, Transcriptional selectors, masters, and combinatorial codes: regulatory principles of neural subtype specification. Wiley Interdiscip Rev Dev Biol 4, 505–528 (2015).

4. D. Apel, M. G. Surette, Bringing order to a complex molecular machine: the assembly of the bacterial flagella. Biochim Biophys Acta 1778, 1851–1858 (2008).

5. H. C. Berg, The rotary motor of bacterial flagella. Annu Rev Biochem 72, 19–54 (2003).

6. D. O. Serra, A. M. Richter, G. Klauck, F. Mika, R. Hengge, Microanatomy at cellular resolution and spatial order of physiological differentiation in a bacterial biofilm. MBio 4, e00103–00113 (2013).

7. R. S. Friedlander, N. Vogel, J. Aizenberg, Role of Flagella in Adhesion of Escherichia coli to Abiotic Surfaces. Langmuir 31, 6137–6144 (2015).

8. J. Haiko, B. Westerlund-Wikstrom, The role of the bacterial flagellum in adhesion and virulence. Biology (Basel) 2, 1242–1267 (2013).

9. N. Q. Balaban, J. Merrin, R. Chait, L. Kowalik, S. Leibler, Bacterial persistence as a phenotypic switch. Science 305, 1622–1625 (2004).

10. I. Keren, D. Shah, A. Spoering, N. Kaldalu, K. Lewis, Specialized persister cells and the mechanism of multidrug tolerance in Escherichia coli. J Bacteriol 186, 8172–8180 (2004).

11. J. E. Gonzalez-Pastor, E. C. Hobbs, R. Losick, Cannibalism by sporulating bacteria. Science 301, 510–513 (2003).

12. H. Maamar, A. Raj, D. Dubnau, Noise in gene expression determines cell fate in Bacillus subtilis. Science 317, 526–529 (2007).

13. A. Chastanet et al., Broadly heterogeneous activation of the master regulator for sporulation in Bacillus subtilis. Proc Natl Acad Sci U S A 107, 8486–8491 (2010).

14. J. W. Veening, W. K. Smits, O. P. Kuipers, Bistability, epigenetics, and bet-hedging in bacteria. Annu Rev Microbiol 62, 193–210 (2008).

15. S. Kalir et al., Ordering genes in a flagella pathway by analysis of expression kinetics from living bacteria. Science 292, 2080–2083 (2001).

16. S. Saini, J. D. Brown, P. D. Aldridge, C. V. Rao, FliZ Is a posttranslational activator of FlhD4C2-dependent flagellar gene expression. J Bacteriol 190, 4979–4988 (2008).

17. D. M. Fitzgerald, R. P. Bonocora, J. T. Wade, Comprehensive mapping of the Escherichia coli flagellar regulatory network. PLoS Genet 10, e1004649 (2014).

18. P. Wang et al., Robust growth of Escherichia coli. Curr Biol 20, 1099–1103 (2010).

19. L. Potvin-Trottier, N. D. Lord, G. Vinnicombe, J. Paulsson, Synchronous long-term oscillations in a synthetic gene circuit. Nature 538, 514–517 (2016).

20. E. Balleza, J. M. Kim, P. Cluzel, Systematic characterization of maturation time of fluorescent proteins in living cells. Nat Methods 15, 47–51 (2018).

21. S. Shin, C. Park, Modulation of flagellar expression in Escherichia coli by acetyl phosphate and the osmoregulator OmpR. J Bacteriol 177, 4696–4702 (1995).

22. D. Lehnen et al., LrhA as a new transcriptional key regulator of flagella, motility and chemotaxis genes in Escherichia coli. Mol Microbiol 45, 521–532 (2002).

23. M. Ko, C. Park, H-NS-Dependent regulation of flagellar synthesis is mediated by a LysR family protein. J Bacteriol 182, 4670–4672 (2000).

24. A. Francez-Charlot et al., RcsCDB His-Asp phosphorelay system negatively regulates the flhDC operon in Escherichia coli. Mol Microbiol 49, 823–832 (2003).

25. N. De Lay, S. Gottesman, A complex network of small non-coding RNAs regulate motility in Escherichia coli. Mol Microbiol 86, 524–538 (2012).

26. J. H. Davis, A. J. Rubin, R. T. Sauer, Design, construction and characterization of a set of insulated bacterial promoters. Nucleic Acids Res 39, 1131–1141 (2011).

27. T. Wada, Y. Hatamoto, K. Kutsukake, Functional and expressional analyses of the anti-FlhD4C2 factor gene ydiV in Escherichia coli. Microbiology 158, 1533–1542 (2012).

28. A. Takaya et al., YdiV: a dual function protein that targets FlhDC for ClpXP-dependent degradation by promoting release of DNA-bound FlhDC complex. Mol Microbiol 83, 1268–1284 (2012).

29. B. Li et al., Structural insight of a concentration-dependent mechanism by which YdiV inhibits Escherichia coli flagellum biogenesis and motility. Nucleic Acids Res 40, 11073-11085 (2012).

30. L. D. Evans, C. Hughes, G. M. Fraser, Building a flagellum outside the bacterial cell. Trends Microbiol 22, 566–572 (2014).

31. F. F. Chevance, K. T. Hughes, Coordinating assembly of a bacterial macromolecular machine. Nat Rev Microbiol 6, 455–465 (2008).

32. M. S. Chadsey, J. E. Karlinsey, K. T. Hughes, The flagellar anti-sigma factor FlgM actively dissociates Salmonella typhimurium sigma28 RNA polymerase holoenzyme. Genes Dev 12, 3123–3136 (1998).

33. J. E. Karlinsey et al., Completion of the hook-basal body complex of the Salmonella typhimurium flagellum is coupled to FlgM secretion and fliC transcription. Mol Microbiol 37, 1220–1231 (2000).

34. G. S. Chilcott, K. T. Hughes, Coupling of flagellar gene expression to flagellar assembly in Salmonella enterica serovar typhimurium and Escherichia coli. Microbiol Mol Biol Rev 64, 694–708 (2000).

35. S. Koirala et al., A nutrient-tunable bistable switch controls motility in Salmonella enterica serovar Typhimurium. MBio 5, e01611–01614 (2014).

36. J. Adler, Chemotaxis in bacteria. Annu Rev Biochem 44, 341–356 (1975).

37. K. Zhao, M. Liu, R. R. Burgess, Adaptation in bacterial flagellar and motility systems: from regulon members to ‘foraging’-like behavior in E. coli. Nucleic Acids Res 35, 4441-4452 (2007).

38. F. C. Neidhardt, R. Curtiss, Escherichia coli and Salmonella : cellular and molecular biology. (ASM Press, Washington, D.C., ed. 2nd, 1996).

39. S. L. Fink, B. T. Cookson, Caspase-1-dependent pore formation during pyroptosis leads to osmotic lysis of infected host macrophages. Cell Microbiol 8, 1812–1825 (2006).

40. E. A. Miao et al., Cytoplasmic flagellin activates caspase-1 and secretion of interleukin 1beta via Ipaf. Nat Immunol 7, 569–575 (2006).

41. C. S. Barker, B. M. Pruss, P. Matsumura, Increased motility of Escherichia coli by insertion sequence element integration into the regulatory region of the flhD operon. J Bacteriol 186, 7529–7537 (2004).

