## Supplementary material for "Stochastic transcriptional pulses orchestrate flagellum biosynthesis in *E. coli*": SM

### Supplementary Information for J.M. Kim et al.

#### A. LIST OF PLASMIDS AND STRAINS

##### Table A1. List of Plasmids

| **Plasmid** | **Source** | **Antibiotic Marker** | **Description** |
| --- | --- | --- | --- |
| pMK4 | This work | Kan | Empty vector with YFP (Venus NB) |
| pMK7 | This work | Amp | Empty vector with CFP (SCFP3A) |
| pMK4-FlgA | This work | Kan | Plasmid containing flgA promoter fused to YFP (Venus NB) |
| pMK4-FlgB | This work | Kan | Plasmid containing flgB promoter fused to YFP (Venus NB) |
| pMK4-FlhB | This work | Kan | Plasmid containing flhB promoter fused to YFP (Venus NB) |
| pMK4-FliA | This work | Kan | Plasmid containing fliA promoter fused to YFP (Venus NB) |
| pMK4-FliD | This work | Kan | Plasmid containing fliD promoter fused to YFP (Venus NB) |
| pMK4-FliE | This work | Kan | Plasmid containing fliE promoter fused to YFP (Venus NB) |
| pMK4-FliF | This work | Kan | Plasmid containing fliF promoter fused to YFP (Venus NB) |
| pMK4-FliL | This work | Kan | Plasmid containing fliL promoter fused to YFP (Venus NB) |
| pMK4-FlgK | This work | Kan | Plasmid containing flgK promoter fused to YFP (Venus NB) |
| pMK4-FlgM | This work | Kan | Plasmid containing flgM promoter fused to YFP (Venus NB) |
| pMK4-FliC | This work | Kan | Plasmid containing fliC promoter fused to YFP (Venus NB) |
| pMK4-MotA | This work | Kan | Plasmid containing motA promoter fused to YFP (Venus NB) |
| pMK4-Tar | This work | Kan | Plasmid containing tar promoter fused to YFP (Venus NB) |
| pMK7-FliF | This work | Amp | Plasmid containing fliF promoter fused to CFP (SCFP3A) |
| pMK7-FliC | This work | Amp | Plasmid containing fliC promoter fused to CFP (SCFP3A) |
| pPro1-YFP | This work | Gent | Plasmid containing Pro1 promoter fused to YFP (Venus NB) |
| pPro2-YFP | This work | Gent | Plasmid containing Pro2 promoter fused to YFP (Venus NB) |
| pPro4-YFP | This work | Gent | Plasmid containing Pro4 promoter fused to YFP (Venus NB) |
| pPro5-YFP | This work | Gent | Plasmid containing Pro5 promoter fused to YFP (Venus NB) |
| pProB-YFP | This work | Gent | Plasmid containing ProB promoter fused to YFP (Venus NB) |
| pSIM5 | Gift of D. Court | Cm | Helper plasmid encoding red recombinase proteins |
| pCP20 | CGSC | Amp, Cm | Helper plasmid encoding FLP recombinase |

##### Table A2. List of Strains

| **Strain** | **Source** | **Description** |
| --- | --- | --- |
| MG1655 (CGSC 6300) | CGSC | Background Strain |
| MG1655+IS5 | This work | See below |
| MGR | This work | MG1655 IntS::ZeoR-P_RNA1_-mCherry |
| MGR-E98K | This work | MGR *motA(E98K)* |
| MGR-E98K FliF FlhD (“E98KFD”) | This work | MGR-E98K galK::AmpR-P_fliF_-CFP flhDC-YFP (see below) |
| MGR-E98K FliF FlgA | This work | MGR-E98K galK::AmpR-P_fliF_-CFP attB::KmR-P_flgA_-YFP |
| MGR-E98K FliF FlgB | This work | MGR-E98K galK::AmpR-P_fliF_-CFP attB::KmR-P_flgB_-YFP |
| MGR-E98K FliF FlhB | This work | MGR-E98K galK::AmpR-P_fliF_-CFP attB::KmR-P_flhB_-YFP |
| MGR-E98K FliF FliA | This work | MGR-E98K galK::AmpR-P_fliF_-CFP attB::KmR-P_fliA_-YFP |
| MGR-E98K FliF FliD | This work | MGR-E98K galK::AmpR-P_fliF_-CFP attB::KmR-P_fliD_-YFP |
| MGR-E98K FliF FliE | This work | MGR-E98K galK::AmpR-P_fliF_-CFP attB::KmR-P_fliE_-YFP |
| MGR-E98K FliF FliF | This work | MGR-E98K galK::AmpR-P_fliF_-CFP attB::KmR-P_fliF_-YFP |
| MGR-E98K FliF FliL | This work | MGR-E98K galK::AmpR-P_fliF_-CFP attB::KmR-P_fliL_-YFP |
| MGR-E98K FliF FliC (“E98KFC”) | This work | MGR-E98K galK::AmpR-P_fliF_-CFP attB::KmR-P_fliC_-YFP |
| MGR-E98K FliC FlgK | This work | MGR-E98K galK::AmpR-P_fliC_-CFP attB::KmR-P_flgK_-YFP |
| MGR-E98K FliC FlgM | This work | MGR-E98K galK::AmpR-P_fliC_-CFP attB::KmR-P_flgM_-YFP |
| MGR-E98K FliC FliC | This work | MGR-E98K galK::AmpR-P_fliC_-CFP attB::KmR-P_fliC_-YFP |
| MGR-E98K FliC MotA | This work | MGR-E98K galK::AmpR-P_fliC_-CFP attB::KmR-P_motA_-YFP |
| MGR-E98K FliC Tar | This work | MGR-E98K galK::AmpR-P_fliC_-CFP attB::KmR-P_tar_-YFP |
| E98KFD Pro1 | This work | MGR-E98K galK::AmpR-P_fliF_-CFP flhDC-YFP P_flhD_::GentR-Pro1 |
| E98KFD Pro2 | This work | MGR-E98K galK::AmpR-P_fliF_-CFP flhDC-YFP P_flhD_::GentR-Pro2 |
| E98KFD Pro4 | This work | MGR-E98K galK::AmpR-P_fliF_-CFP flhDC-YFP P_flhD_::GentR-Pro4 |
| E98KFD Pro5 | This work | MGR-E98K galK::AmpR-P_fliF_-CFP flhDC-YFP P_flhD_::GentR-Pro5 |
| E98KFD ProB | This work | MGR-E98K galK::AmpR-P_fliF_-CFP flhDC-YFP P_flhD_::GentR-ProB |
| E98KFD ΔydiV | This work | MGR-E98K galK::AmpR-P_fliF_-CFP flhDC-YFP ydiV::FRT |
| E98KFC Pro4 ΔflgM | This work | MGR-E98K galK::AmpR-P_fliF_-CFP flhDC-YFP P_flhD_::GentR-Pro4 flgM::FRT |
| E98KFD Pro1 ΔydiV | This work | MGR-E98K galK::AmpR-P_fliF_-CFP flhDC-YFP P_flhD_::GentR-Pro1 ydiV::FRT |
| E98KFD Pro2 ΔydiV | This work | MGR-E98K galK::AmpR-P_fliF_-CFP flhDC-YFP P_flhD_::GentR-Pro2 ydiV::FRT |
| E98KFD Pro4 ΔydiV | This work | MGR-E98K galK::AmpR-P_fliF_-CFP flhDC-YFP P_flhD_::GentR-Pro4 ydiV::FRT |
| E98KFD Pro5 ΔydiV | This work | MGR-E98K galK::AmpR-P_fliF_-CFP flhDC-YFP P_flhD_::GentR-Pro5 ydiV::FRT |
| E98KFD ProB ΔydiV | This work | MGR-E98K galK::AmpR-P_fliF_-CFP flhDC-YFP P_flhD_::GentR-ProB ydiV::FRT |
| E98KFC+IS5 | This work | MGR-E98K+IS5 galK::AmpR-P_fliF_-CFP attB::KmR-P_fliC_-YFP |

#### B. STRAIN CONSTRUCTION

##### a. Background Strain

The background strain for this work was MG1655 (Coli Genetic Stock Center (CGSC) #6300). Two stocks of MG1655 are commonly available: MG1655 (seq) (CGSC #7740) which derives from the subculture used by the Blattner lab for the complete genomic sequencing of *E. coli* harbors an IS1 element in the regulatory region of FlhDC (*1*). By contrast, CGSC #6300, the original isolate of MG1655 submitted by the Blattner lab, does not have this insertion element sequence. For a discussion of the impact of this insertion element, see the section below “Impact of Insertion Element Mutation on Flagellar Transcription”.

##### b. Chromosomal Engineering

Strains used in this study were constructed using red recombination (*2*). Parental strains were transformed with the helper plasmid pSIM5 and red recombination was performed using standard protocols (*3*). During strain construction, cells were grown at 30°C to ensure maintenance of pSIM5, which has a temperature sensitive origin. All strains were cured of the helper plasmid prior to the experiment.

##### c. “Scarless” Chromosomal Engineering

To generate point-mutations or insert sequences without selective markers, we used a “scarless” chromosomal engineering technique—i.e. a genome editing strategy that eliminates extraneous sequences such as antibiotic resistance cassettes (*4*). We modified a dual selection/counter-selection cassette in pKD45 (*5*) consisting of a kanamycin resistance marker and a toxin ccdB driven by a rhamnose inducible promoter (P_RhaB_). This original cassette required counter-selection to be done on minimal M9 plates containing rhamnose because presence of other carbon sources allowed cells to avoid activation of the rhamnose promoter and escape counter selection. Cell growth was extremely slow on these plates, requiring almost two full days of incubation before colonies became visible. To overcome this limitation, we replaced the rhamnose inducible promoter with the arabinose inducible P_araB_ using standard molecular biology techniques and isothermal assembly (*6*) using the NEB HiFi DNA Assembly Master Mix (New England Biolabs). The resulting *kmR-araBp-ccdB* (“KAC”) cassette allowed for efficient counter-selection on LB plates containing 10% arabinose. We also constructed a variant of this cassette with the gentamicin resistance (*gmR-araBp-ccdB* or “GAC”).

##### d. Insertion of Constitutive RFP Marker

For our experiments, the background strain was first engineered to constitutively express mCherry. This modification improved identification of cells in flow cytometry experiments and facilitated segmentation of cells in time-lapse experiments. An expression cassette containing (i) a Zeocin resistance marker, (ii) a RNA-I promoter from ColE1 fused to a coding sequence for mCherry and (iii) two 100 bp long flanking sequences homologous to the IntS gene was ordered from IDT as a gBlock®. This linear fragment was chromosomally inserted into the IntS locus using the chromosomal engineering techniques described above. The insertion did not affect the growth rate under our experimental conditions or flagellar promoter activity as determined by flow cytometry. The strain harboring the constitutive red marker was designated “MGR” (MG1655 “Red”).

##### e. Point Mutation of MotA

Motile cells can swim out of the channels in the “mother machine”. To prevent the loss of cells expressing flagella from the channels, we engineered the point mutant MotA(E98K) (*7*) directly in the chromosome using the KAC cassette described above. This point mutation prevents MotA from powering the rotation of the flagellar motor. We confirmed via flow cytometry that this point mutation did not affect flagellar gene expression. For all time-lapse studies in the mother machine, a MGR strain harboring the MotAE98K mutation (henceforth termed “E98K”) was used as the background strain.

##### f. Construction of Transcriptional Reporters

To generate the library of transcriptional reporters, we first generated the cloning vectors pMK4 and pMK7 which were generated from pUA66 (*8*). In pMK4, the GFP coding sequence and ribosomal binding site (RBS) were replaced by the RBS of Gene 10 in T7 phage (“T7 RBS”) and Venus NB (“VenNB”) (*9*). In pMK7, the GFP coding sequence and RBS were replaced by the T7 RBS and SCFP3A (*9*). In both cases, we empirically optimized the coding sequence of VenNB and SCFP3A for high translation rate via random mutagenesis. In pMK7, the kanamycin resistance cassette and the lambda T0 terminator were also replaced with an ampicillin resistance cassette and a high efficiency terminator from the ilvGEDA regulatory region (*10*). In addition to these changes, the original restriction cloning site was replaced with a region flanked by two BsaI recognition sites to allow for insertion of promoter sequences via Golden Gate assembly (*11, 12*). Both pMK4 and pMK7 were constructed by assembling PCR amplified cassettes into a circular plasmid via isothermal assembly using the NEB HiFi DNA Assembly Master Mix (New England Biolabs).

For each transcriptional reporter, we amplified via PCR a region of the chromosome encompassing the flagellar promoter of interest as well as >30bp of the coding sequences flanking the promoter. This strategy was implemented in case any previously unknown regulator binding sites existed in that region. The PCR primers were designed to contain overhangs that contained BsaI sites as well as target sequences to pMK4 or pMK7 that allowed for the assembly of the PCR product with pMK4 or pMK7 via Golden Gate assembly. The Golden Gate assembly reaction was performed as previously described, using the higher efficiency protocol of cycling between 3min incubation at 37°C and 4 min incubation at 16°C for 25 cycles (*11*).

The transcriptional reporters were inserted into one of two chromosomal loci. The first locus was the lambda attB site in the chromosome. The second locus was the GalK. Both loci have previously been used for the insertion of chromosomal transcriptional reporters (*13-15*). The two sites are approximately 20kb apart, which is sufficiently far to prevent read-through transcription (this read-through is already low due to the strong rrnB T1 terminator at the end of the VenNB cassette) from interfering with the measurements. The two sites, however, are sufficiently close to each other so that variations in chromosomal copy number during replication can safely be ignored. Integration into both sites was performed by amplifying the transcriptional reporter from the plasmid template with primers each containing at least 40bp overhang homologous to the target locus. This PCR fragment was inserted chromosomally using the techniques described above.

##### g. Construction of Class 1 Transcriptional Reporter

Unlike other flagellar promoters, the regulatory region of the Class 1 gene FlhDC is highly complex. Moreover, evidence suggests that transcription from this promoter may be sensitive to long range interactions (*1, 16*). Therefore, we decided to examine transcription of Class 1 genes by directly integrating a cassette composed of the T7 RBS and VenNB into the 3’ UTR of the FlhDC transcript. One complication arises from the fact that the promoter of the neighboring gene MotA resides within the FlhC coding sequence. To avoid interference from this promoter, we replaced the codons of FlhC comprising the MotA promoter with synonymous codons. Both the synonymous mutations and the VenNB insertion was accomplished using the “scarless” technique described above. To confirm that the insertion did not significantly alter downsteam flagellar gene expression, we inserted a Class 2 reporter in this strain and compared the distribution of fluorescence with that of the Class 2 reporter in a wild-type strain (**Fig. S4**).

#### C. GROWTH CONDITIONS

For all experiments, except where otherwise indicated, we used a modified version of the Neidhardt EZ rich media (Teknova), an optically clear rich defined media based on a MOPS buffer (*17*). We replaced the 0.2% glucose in the original formulation with 0.4% glycerol to prevent catabolite repression of flagellar synthesis.

For initial experiments, cells were grown at 34ºC so that the width of the cell better matches the width of our initial microfluidic device. Subsequently, we generated additional microfluidic devices with narrower channels that permitted cell growth at 30ºC which, due to the slower generation time, permitted more fields of view to be sampled in a single experiment. The pulsating dynamics were qualitatively similar at both temperatures. Regardless, for any set of experiments where we compared multiple strains (e.g. pairwise Class II and Class III measurements, wild-type vs mutant, synthetic flhDC expression), we used identical temperature and growth conditions to enable proper quantitative comparison.

#### D. DATA ACQUISITION AND ANALYSIS

##### a. Flow Cytometry

For flow cytometry experiments, an overnight culture was diluted at least 1:3000 into 500µl of fresh growth media. Cells were incubated at 30°C with shaking at 250rpm for at least 6 hours prior to measurement. At this point, the cell density was typically around or below OD=0.1.

Cells were analyzed on a BD Fortessa flow cytometer (Becton Dickinson). The side-scatter (SSC-A) profile was used to first discriminate cells from background particles. Where possible, the constitutive red fluorescence from the mCherry marker was used as a second gate. The raw flow cytometry data was imported into MATLAB (Mathworks) and analyzed via custom software (**Fig. S3, S4, S7-S9**).

##### b. Time-Lapse Experiments in the Microfluidic Device

###### i) Microfluidic master fabrication

For our initial experiments, we used an epoxy replica of the “mother machine” described in Potvin-Trottier et al (*18*) (**Fig. S1**, Design A). The replica mold was a generous gift from Dr. Matthew Cabeen (Harvard University). Subsequently, we generated custom SU-8 molds to build channels that better matched the typical cell width in our growth conditions. New designs for the microfluidic device were created in AutoCAD; generally, it consists of two layers, one for the cell channels and a second one for the feeding channels. The cell channels were 1.1μm wide and 25μm long. The edges of the channels were smoothed out to reduce the halo that appears during phase contrast imaging. We also added a trough at the boundary where the cell channels meet the feeding channel so that the phase contrast halo from the feeding channel does not affect the imaging of the cells inside the channel (**Fig. S1**, Design B). The feeding channels were 8.1mm long and 100μm wide.

Fabrication of the master mold was carried out using standard UV photolithography in a clean room environment at the Center for Nanoscale Systems at Harvard University. We modified the fabrication procedure from the method described in (*18*) by exposing the SU-8 using a Heidelberg MLA150 Maskless Aligner (Heidelberg Instruments). The MLA150 enabled us to directly “print” our AutoCAD designs without a mask and often resulted in more accurate printing of smaller features.

To print the microfluidic device master, we used the following protocol. The spin coating parameters shown below are written using the abbreviation: speed (rpm)/acceleration (rpm/sec)/time (sec):

1) First layer: cell channels.
   a. Place a 3″ wafer at a spinner and rinse it by adding acetone and isopropyl alcohol (IPA) while it is spinning.
   b. Let the wafer dry for 15 minutes on a hotplate at 200°C.
   c. Let the wafer cool down for a few minutes and place it on a spin chuck.
   d. Slowly pour SU-8 2002 until it covers ~2/3 of the wafer. Here, avoid any bubbles in the resist since even small bubbles can distort the cell channel.
   e. Spin the wafer using the program: Step 1: 500/100/10, Step 2: 3500/300/60.
   f. Bake the wafer for 1 min at 65°C, 1 min at 95°C, 1 min at 65°C.
   g. Expose the wafer with the cell channel design using the MLA150 with a dosage of 2500 mJ/cm^2^. In our cell channel design, we also include cross shaped marks which will serve as alignment marks during the exposure of the feeding channel layer.
   h. After exposure in the MLA, bake the wafer for 1 min at 65°C, 1 min 95°C and 30 seconds at 65°C.
   i. Gently immerse the wafer in a SU-8 developer.
   j. Bake the wafer for 15 minutes on a hot plate at 150°C (“hard bake” step).
   k. Measure the channel height using a profilometer. The expected high is ~1.2μm.
2) Second layer: feeding channels.
   a. Place the wafer with the cell channels on a spin chuck.
   b. Slowly pour SU-8 2010 photoresist on the wafer covering ~2/3 of its area.
   c. Spin the wafer using the program: Step 1: 500/100/10. Step 2: 3000/300/60.
   d. Bake it using hot plates for 1 min at 65°C, 2 min 95°C and 1 min at 65°C.
   e. Use cotton swabs soaked with propylene-glycol-methyl-ether-acetate (PGMA) to wipe SU-8 off from the region of the wafer where the alignment crosses are printed.
   f. Bake the wafer at 65°C for 1 minute.
   g. Load the feeding channel design into the MLA. Place the wafer in the MLA and align the wafer by identifying the crosses using the cameras of the MLA. Once alignment is complete, expose with a dosage of 4500 mJ/cm^2^ and focus offset (“defoc”) -2.
   h. Once the design is exposed, bake the wafer at 65°C for 1 min, 95°C for 4 min and 65°C for 1 min.
   i. Immerse the wafer in a container with PGMA and shake it very slowly for 1 min.
   j. Rinse the wafer with IPA to remove the remaining SU-8.
   k. Let the wafer hard bake at 150ºC for 15 minutes.
   l. Measure the feeding channel height using the profilometer. The expected height is ~11μm.

###### ii) Microfluidic device fabrication

To prepare a new microfluidic device out of the mold, dimethyl siloxane monomer (Sylgard 184, Dow Corning) was mixed with the curing agent at a 10:1 ratio, degassed and poured over the mold. This mixture was degassed for an additional 1 hour and cured overnight at 65°C before being removed from the mold. Individual devices were first cut from the cured PDMS. Subsequently, inlets and outlets for each flow channel (“lane”) were created using a 0.75 mm biopsy punch. The device was treated with oxygen plasma in a plasma cleaner along with a 25mm×40mm No. 1.5 coverglass (VWR) for 15s at 30W and oxygen pressure 200mTorr. Following plasma treatment, the device and glass were bonded and incubated at least 1 hour at 65°C prior to use.

###### iii) Cell and growth media preparation

For time-lapse experiments, we added a passivating agent Pluronic F-108 (Sigma Aldrich) at a final concentration of 0.85g/L to the growth media. *E. coli* strains were first grown overnight in media without the passivating agent. Cells were diluted the morning of the experiment 1:100 fold in fresh media, containing the passivating agent and allowed to grow to late exponential or early stationary phase. The reduced cell size at this growth phase improved the efficiency with which cells loaded in the device.

The cell culture was loaded into the inlet of the device by pipetting. The device was then centrifuged on a custom adaptor fit into a standard table-top centrifuge at 6000×g for 10 mins. The inlets were connected to syringes filled with the growth media (+passivating agent) via Tygon® tubing (VWR, ID 0.02″×OD 0.06″). The flow-through was collected from the outlet via a second Tygon® tubing into an empty beaker. The growth media was first pumped at a rate of 35µl/min for at least 1 hour to allow for the inlets and outlets to be cleared. Afterwards, the flow rate was reduced to 4-5µl/min for the duration of the experiment.

###### iv) Imaging protocol

The cells were allowed to adapt to growth in the device for at least 2 additional hours before imaging. Time-lapse images in experiments performed at 34ºC were acquired on a Nikon Eclipse Ti inverted microscope equipped with a 60× Plan Apo oil objective (numerical aperture (NA) 1.4, Nikon), an Orca R2 CCD camera (Hamamatsu), an automated xy-stage (Ludl) and a SOLA light engine LED excitation source (Lumencor). The microscope was also surrounded by a temperature controlled enclosure. The following filter sets were used for acquisition: YFP (Semrock YFP-2427A), CFP (Semrock CFP-2432A), and RFP (Semrock mCherry-A). Automated time-lapse acquisition was controlled using custom MATLAB 2011a (Mathworks) scripts interfacing with μManager 1.4.

Time-lapse images in subsequent experiments at 30ºC were acquired on a Zeiss Axiovert 200M microscope equipped with a Plan-Apochromat 40x/1.3 Oil Ph3 Objective and a CCD camera (Hamamatsu C4742-98- 24ERG). For fluorescence excitation we used an LED illumination source, SOLA SE II (Lumencor). Filters with the following specifications were used: for YFP (Ex. 500/24, Di. 520, Em.542/27), CFP (Ex. 438/24, Di. 458, Em. 483/32) and RFP (Ex. 586/20, Di. 605, Em. 647/57). All filters were from Semrock. The variation in fluorescence intensity illumination across the field-of-view was less than 10% in all channels. The microscope setup was controlled using custom software on MATLAB 2013a (Mathworks) interfacing with µManager 1.4.

###### v) Time-lapse acquisition parameters

In the Nikon microscope, images were taken every 10 minutes and focal drift was corrected via the Nikon PerfectFocus system and periodically recalibrated using z-stacks on a sacrificial position. Images in the RFP (the cell segmentation marker) were acquired at full camera resolution (1344×1024 pixels) to improve segmentation while CFP and YFP images were acquired using 2×2 binning to reduce measurement noise. Short exposure times (typically 200-300ms) and low illumination intensities (<30% of maximum illumination power) were used to minimize the effects of photobleaching.

In the Zeiss microscope, automated time-lapse acquisition was controlled using custom MATLAB (Mathworks) scripts interfacing with μManager. Images were acquired every 5 minutes. Focal drift was corrected at each acquisition step via a custom autofocus routine which acquires phase contrast images at planes above and below the previously determine optimal autofocus plane and estimates the image plane with maximal contrast. Once the new focal plane was determined, images in the YFP, CFP and RFP channels were acquired at full camera resolution (1344×1024 pixels). Again, short exposure times (typically 200-300ms) and low illumination intensities (<15% of maximum illumination power) were used to minimize the effects of photobleaching.

##### c. Tracking Software

We developed custom software in MATLAB to analyze our time-lapse movies. Using this software, we could identify and track mother cells lineages with only occasional manual supervision. The software implements the following steps:

###### i) Identification of individual cells in a single frame

For each frame, individual cells were identified and segmented using custom software implementing previously described algorithms (*18-20*). We used RFP fluorescence as the “reference” image channel for segmentation. Parameters of the segmentation algorithm were optimized for our typical image conditions.

###### ii) Correction for shift between fluorescence channels

Once the “masks” defining individual cells were determined, images from the YFP and CFP data channels were aligned to the RFP segmentation to correct for any mis-registration between the data channels. The horizontal shift between the reference RFP channel and the data channel (YFP & CFP) was computed by taking the horizontal projection of each channel and computing the cross-correlation function using the *xcov* function in MATLAB: the shift between fluorescence channels was estimated by determining the lag value where the maximum cross-correlation coefficient occurs. The shift between the fluorescence channels is usually very small (1-2 pixels). However, at large lag values, computationally estimated cross-correlations can occasionally generate large (spurious) correlation values. To avoid this problem, we limited the range of lags being considered to ±5 pixels. Similarly, we estimated the vertical shift using a cross-correlation analysis of the vertical projections for each fluorescence channel.

###### iii) Tracking of the mother cell lineage

In most cases, the cell at the bottom of the growth channel (i.e. the “mother cell”) remained virtually in the same position from frame-to-frame. Thus, the mother cell lineage could be easily identified by following cells closest to the “mother cell” in the previous frame. In some cases, there was a small drift in the field-of-view over the course of the experiment. However, because this drift was relatively small, it could easily be corrected by re-aligning each frame so that the mother cell remained in the center of the field of view. Potential “cell divisions” events were first identified by sudden decreases in cell area, i.e. if a cell’s area dropped to less than 60% of its current value in the next frame. Manual review revealed that most lineages were properly constructed. Occasionally, however, due to mis-segmentation, the cell area increased or decreased precipitously. In most cases, we corrected these errors manually. However, we also implemented a “voting mechanism” similar to the algorithm described in (*21*) to automatically correct these errors. We nonetheless note that measurements of mean fluorescence are very robust to these segmentation errors: erroneous over-segmentation results in two symmetric “half-cells” with the same mean fluorescence as the original full size cell while erroneous under-segmentation results in fluorescence of two daughter cells being averaged together which, due to the shared parental history, tends to be relatively similar (see further discussion below in “Time-lapse Data Analysis”).

Each lineage was tracked to the end of the experiment or until the mother cells were “lost” from the channels. For simplicity, only lineages that survived and continued to divide to the end of the experiment were analyzed. Also, in rare occasions, mother cells became filamentous or stopped growing altogether. These lineages were also discarded from analysis.

###### iv) Measurement of fluorescence and cell length

Once the cells corresponding to the mother cell lineage were identified in each frame, we extracted the mean fluorescence and cell length measurements. Mean fluorescence was computed by collecting and averaging the pixel values of the fluorescence image that lie within the mask corresponding to the mother cell. We computed the mean fluorescence for all three (YFP, CFP and RFP) fluorescence channels. For each mother cell, we also estimated the approximate cell length: due to the vertical (or near-vertical) orientation of the almost all mother cells, the distance between the top- and bottom-most pixels of the mother cell was taken to be reasonable estimates of the cell length.

##### d. Time-lapse Data Analysis

###### i) Estimation of promoter activity

The activity of a promoter driving a transcript expressing a fluorescent protein can be estimated by the amount of new fluorescent protein produced between two time points). The fluorescent proteins used in our experiments are generally very stable and undergo negligible degradation. Under such conditions:

$$F_{total}\left( t \right)\cong F_{total}\left( t-1 \right)+P(t)$$

where F_total_ is the total fluorescence of the cell and P(t) is the amount of new fluorescence produced between *t-1* and *t*. Thus, in principle, we could estimate *P* from the change is total fluorescence within the cell between two frames. However, estimation of the total fluorescence is sensitive to exact segmentation of each individual cell. Therefore, we instead opted to use methods of promoter activity estimation based on changes in the mean fluorescence (i.e. the average pixel intensity) in the cell as previously described in (*18, 22*)**.** Briefly, if the total area of the cell is A(t) and the average pixel intensity is *C(t)*, then $F_{total}=A\left( t \right)C(t)$. It follows that $\frac{dF_{total}}{dt}=C\frac{dA}{dt}+A\frac{dC}{dt}$ which can be rearranged to give:

$$\frac{1}{A}\frac{dF_{total}}{dt}=C\frac{1}{A}\frac{dA}{dt}+\frac{dC}{dt}$$

We chose to use the left term $\frac{1}{A}\frac{dF_{total}}{dt}$, i.e. the “cell-size normalized” production rate, as a proxy for promoter activity. Because the transcriptional pulses we observed were extremely large and lasted multiple cell generations, this normalization had a negligible effect while simplifying our promoter activity estimation (**Fig. S5**). Operationally, we estimated $\frac{1}{A}\frac{dA}{dt}$, i.e. the relative growth rate of the cell, by taking the log ratio of the initial and final area of the cell for each cell division. We estimated $\frac{dC}{dt}$ by smoothing our fluorescence traces with a Savitzky-Golay filter and taking the numerical derivative.

###### ii) Estimation of “on” and “off” states

To estimate the duration of “on” and “off” states, we first applied a mild Savitzky-Golay filter to our promoter traces. We then defined a heuristic threshold to identify the “pulse on” and “pulse off” states for each promoter. First, using cells without any fluorophores, we estimated the background “promoter activity” of *E. coli* under our experimental conditions which arises due to autofluorescence. From this measurement, we obtained the mean and standard deviations of autofluorescence-associated background activity. We defined the mean + 2×standard deviation of this background activity as our “low detection threshold”. When we applied this threshold to our Class II and Class III promoter activity traces, we discovered that this threshold was too sensitive and that we detected many small “bursts” of transcription lasting under 10 minutes. We hypothesized that those bursts might be “leaky” transcripts that occur even during inactive states. Therefore, collected all short bursts, lasting 10 minutes or less and computed the mean and standard deviation of those bursts. This process allowed us to define a second threshold as the + 2×standard deviation of “leaky transcript” activity. Time periods when the promoter activity was continuously above this threshold was defined as “on” periods. Similarly, “off” periods were defined as contiguous time periods with promoter values below this threshold.

###### iii) Correlation analysis

To compute the average autocorrelation function for a given promoter, we first determined the autocorrelation function of each individual promoter activity trace (each corresponding to a single continuous mother cell lineage) using the *xcov* function in MATLAB. We then averaged the autocorrelation functions for all the traces.

The Pearson correlation coefficients between two promoter activity traces (or between YFP fluorescence and CFP promoter activity) was computed using the *corr* function in MATLAB. The mean correlation coefficient was then obtained by averaging the correlation coefficients from independent lineages.

###### iv) Input-output relationship between promoters across different classes

To determine the input-output relationship between Class II and Class III, we divided the Class II reporter fluorescence into 8 logarithmically-spaced bins that spanned the minimum and maximum Class II fluorescence values measured. For each bin we computed the mean input (i.e. mean Class II fluorescence) against the mean of the corresponding output (i.e. Class III promoter activity).

We used a similar approach to determine the input-output relationship between Class I and Class II in strains where Class I was driven by different synthetic promoters. However, unlike the Class II reporter, the Class I reporter fluorescence has a much smaller variance (see **Fig. S6**) and therefore it was more difficult to distinguish between periods of sustained high fluorescence and short random fluctuations, possibly due to measurement or reporter “noise”. To overcome this effect, we binned our time-lapse data into 2 hour windows (corresponding to ~2 cell divisions) and averaged the Class I fluorescence and Class II promoter activity for each time bin. This step effectively allowed to determine whether the Class I fluorescence and Class II promoter activity was “consistently” high over a given 2 hour time period. We then divided the resulting time-binned Class I reporter values into 5 logarithmically-spaced amplitude-bins that spanned the minimum and maximum Class I fluorescence values measured. For each bin, we computed the mean input (i.e. mean Class I fluorescence) against the mean of the corresponding output (i.e. Class II promoter activity).

#### E. IMPACT OF INSERTION ELEMENT MUTATION ON FLAGELLAR TRANSCRIPTION

The heterogeneity in flagellar gene expression appears to contradict previous physiological observations that mid-exponential phase cells have on average 2-3 flagella each (*23-25*). However, as noted before, traditional strains used in chemotaxis studies (such as RP437 or W3110) have been shown to harbor in insertion element in the regulatory region of *flhDC* (*1*). The experimental strain MG1655 used here, however, has the wild-type regulatory sequence without the insertion element mutation. To determine whether the apparent difference in flagellar gene regulation might be specifically due to the insertion element mutation, we replaced the native FlhDC regulatory region with a homologous region from W3110 which carries an IS5 insertion. This “IS5” strain of MG1655 began expressing flagellin homogenously at a high level (**Fig. S8**). This result supports our hypothesis that the pulsing behavior may have been previously obscured in many common *E. coli* strains that harbor the insertion element.

#### F. IMPACT OF CLASS II AND CLASS III GENES ON CLASS II PULSES

In *Salmonella,* two Class II genes were shown to participate in feedback regulation of FlhDC. The Class II gene product FliZ in *Salmonella* acts as a transcriptional repressor for YdiV (*26*). Hence, expression of FliZ leads to increased activation of FlhDC. A second Class II protein FliT was found to directly bind the FlhC subunit of FlhDC and prevent the complex from associating with DNA (*27, 28*). However, FliT was unable to interact with FlhDC that was pre-bound to DNA (*27*).

By contrast, in *E. coli,* we found that neither FliZ nor FliT had a significant effect on heterogeneous Class 2 gene expression (**Fig. S9**). For FliZ, we note that the 5’ untranslated region of YdiV in *Salmonella* and *E coli* show considerable divergence: transcription of YdiV in *Salmonella* is thought to be regulated by the neighboring nlpC gene promoter, while its *E. coli* counterpart is thought to be expressed from its own promoter within this region (*29*). Thus, we hypothesize that the FliZ-YdiV positive feedback may be a *Salmonella* specific feature.

We also investigated whether FliA, the sigma factor responsible for Class III activation, could potentially affect Class II pulses via feedback interaction. However, deletion of FliA did not substantially alter the distribution of Class II gene expression (**Fig. S9**). Since Class III genes cannot be expressed in ∆fliA, by extension we also believe that Class III gene products do not influence Class II pulses.

Finally, we investigated whether the flagellar basal body complex could potentially influence Class II pulses. To test this hypothesis, we deleted the class II protein FliF which was previously been shown to be necessary for the formation of flagellar basal bodies (*30*). Again, we observed no significant difference in Class II gene expression in this strain (**Fig. S9**) suggesting that Class II pulses occur independently of basal body assembly/inheritance.

**References**

1.	C. S. Barker, B. M. Pruss, P. Matsumura, Increased motility of Escherichia coli by insertion sequence element integration into the regulatory region of the flhD operon. *J Bacteriol* **186**, 7529-7537 (2004).

2.	N. G. Copeland, N. A. Jenkins, D. L. Court, Recombineering: a powerful new tool for mouse functional genomics. *Nat Rev Genet* **2**, 769-779 (2001).

3.	S. K. Sharan, L. C. Thomason, S. G. Kuznetsov, D. L. Court, Recombineering: a homologous recombination-based method of genetic engineering. *Nat Protoc* **4**, 206-223 (2009).

4.	X. T. Li, L. C. Thomason, J. A. Sawitzke, N. Costantino, D. L. Court, Positive and negative selection using the tetA-sacB cassette: recombineering and P1 transduction in Escherichia coli. *Nucleic Acids Res* **41**, e204 (2013).

5.	T. Kolmsee, R. Hengge, Rare codons play a positive role in the expression of the stationary phase sigma factor RpoS (sigma(S)) in Escherichia coli. *RNA Biol* **8**, 913-921 (2011).

6.	D. G. Gibson *et al.*, Enzymatic assembly of DNA molecules up to several hundred kilobases. *Nat Methods* **6**, 343-345 (2009).

7.	Y. V. Morimoto, S. Nakamura, N. Kami-ike, K. Namba, T. Minamino, Charged residues in the cytoplasmic loop of MotA are required for stator assembly into the bacterial flagellar motor. *Mol Microbiol* **78**, 1117-1129 (2010).

8.	A. Zaslaver *et al.*, A comprehensive library of fluorescent transcriptional reporters for Escherichia coli. *Nat Methods* **3**, 623-628 (2006).

9.	E. Balleza, J. M. Kim, P. Cluzel, Systematic characterization of maturation time of fluorescent proteins in living cells. *Nat Methods* **15**, 47-51 (2018).

10.	G. Cambray *et al.*, Measurement and modeling of intrinsic transcription terminators. *Nucleic Acids Res* **41**, 5139-5148 (2013).

11.	C. Engler, R. Kandzia, S. Marillonnet, A one pot, one step, precision cloning method with high throughput capability. *PLoS One* **3**, e3647 (2008).

12.	C. Engler, R. Gruetzner, R. Kandzia, S. Marillonnet, Golden gate shuffling: a one-pot DNA shuffling method based on type IIs restriction enzymes. *PLoS One* **4**, e5553 (2009).

13.	K. A. Datsenko, B. L. Wanner, One-step inactivation of chromosomal genes in Escherichia coli K-12 using PCR products. *Proc Natl Acad Sci U S A* **97**, 6640-6645 (2000).

14.	M. J. Dunlop, R. S. Cox, 3rd, J. H. Levine, R. M. Murray, M. B. Elowitz, Regulatory activity revealed by dynamic correlations in gene expression noise. *Nat Genet* **40**, 1493-1498 (2008).

15.	M. B. Elowitz, A. J. Levine, E. D. Siggia, P. S. Swain, Stochastic gene expression in a single cell. *Science* **297**, 1183-1186 (2002).

16.	C. Lee, C. Park, Mutations upregulating the flhDC operon of Escherichia coli K-12. *J Microbiol* **51**, 140-144 (2013).

17.	F. C. Neidhardt, P. L. Bloch, D. F. Smith, Culture medium for enterobacteria. *J Bacteriol* **119**, 736-747 (1974).

18.	L. Potvin-Trottier, N. D. Lord, G. Vinnicombe, J. Paulsson, Synchronous long-term oscillations in a synthetic gene circuit. *Nature* **538**, 514-517 (2016).

19.	T. M. Norman, N. D. Lord, J. Paulsson, R. Losick, Memory and modularity in cell-fate decision making. *Nature* **503**, 481-486 (2013).

20.	S. Uphoff *et al.*, Stochastic activation of a DNA damage response causes cell-to-cell mutation rate variation. *Science* **351**, 1094-1097 (2016).

21.	Y. Yang, X. Song, A. Lindner, Time-lapse microscopy and image analysis of Escherichia coli cells in mother machines. *Methods in Microbiology* **43**, 49-68 (2016).

22.	J. H. Levine, M. E. Fontes, J. Dworkin, M. B. Elowitz, Pulsed feedback defers cellular differentiation. *PLoS Biol* **10**, e1001252 (2012).

23.	H. Makinoshima *et al.*, Growth phase-coupled alterations in cell structure and function of Escherichia coli. *J Bacteriol* **185**, 1338-1345 (2003).

24.	H. Tang, D. F. Blair, Regulated underexpression of the FliM protein of Escherichia coli and evidence for a location in the flagellar motor distinct from the MotA/MotB torque generators. *J Bacteriol* **177**, 3485-3495 (1995).

25.	L. Turner, W. S. Ryu, H. C. Berg, Real-time imaging of fluorescent flagellar filaments. *J Bacteriol* **182**, 2793-2801 (2000).

26.	T. Wada, Y. Tanabe, K. Kutsukake, FliZ acts as a repressor of the ydiV gene, which encodes an anti-FlhD4C2 factor of the flagellar regulon in Salmonella enterica serovar typhimurium. *J Bacteriol* **193**, 5191-5198 (2011).

27.	Y. Sato, A. Takaya, C. Mouslim, K. T. Hughes, T. Yamamoto, FliT selectively enhances proteolysis of FlhC subunit in FlhD4C2 complex by an ATP-dependent protease, ClpXP. *J Biol Chem* **289**, 33001-33011 (2014).

28.	S. Yamamoto, K. Kutsukake, FliT acts as an anti-FlhD2C2 factor in the transcriptional control of the flagellar regulon in Salmonella enterica serovar typhimurium. *J Bacteriol* **188**, 6703-6708 (2006).

29.	T. Wada, Y. Hatamoto, K. Kutsukake, Functional and expressional analyses of the anti-FlhD4C2 factor gene ydiV in Escherichia coli. *Microbiology* **158**, 1533-1542 (2012).

30.	H. Li, V. Sourjik, Assembly and stability of flagellar motor in Escherichia coli. *Mol Microbiol* **80**, 886-899 (2011).

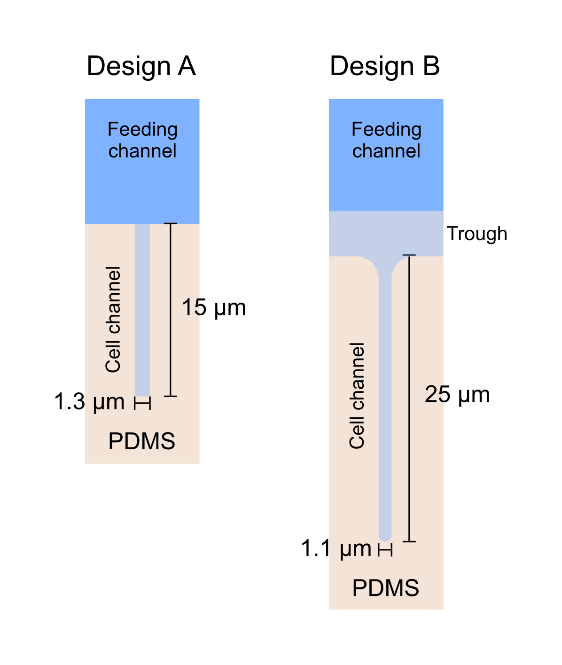

**Fig. S1. Schematic of the microfluidic device.** (Left) Design A used for initial experiments. This design accommodates ~3-4 cells per cell channel. (Right) Design B incorporates several modifications. The cell channels are thinner (1.1μm vs 1.3μm) and extended to 25μm long to accommodate ~6-8 cells. The narrower channels are better suited for maintaining cells in a vertical orientation under our growth conditions. The edges of the channels are rounded which reduces the halo effect during phase contrast microscopy. Finally, a shallow trough is placed between the cell channel and the feeding channel so that the phase contrast halo from the feeding channel does not interfere with imaging of the cells within the cell channel.

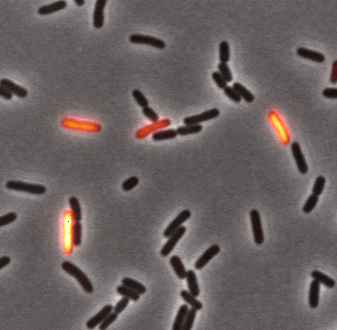

**Fig. S2. Flagellar gene expression is heterogeneous when cells are grown in liquid suspension.** Example image of cells with a fliCp-YFP reporter grown in liquid culture. Shown is an overlay of phase contrast and YFP fluorescence (orange) images. Cells were grown in liquid culture at 30ºC in the modified Neidhardt EZ rich media (see section “Growth Conditions” from above) with shaking at 250rpm until OD=0.5. 1μl of the culture was then deposited onto an agarose pad (2% w/v, low melting point agarose BP165, Fisher Scientific) and a coverslip was placed on top to allow imaging under the microscope.

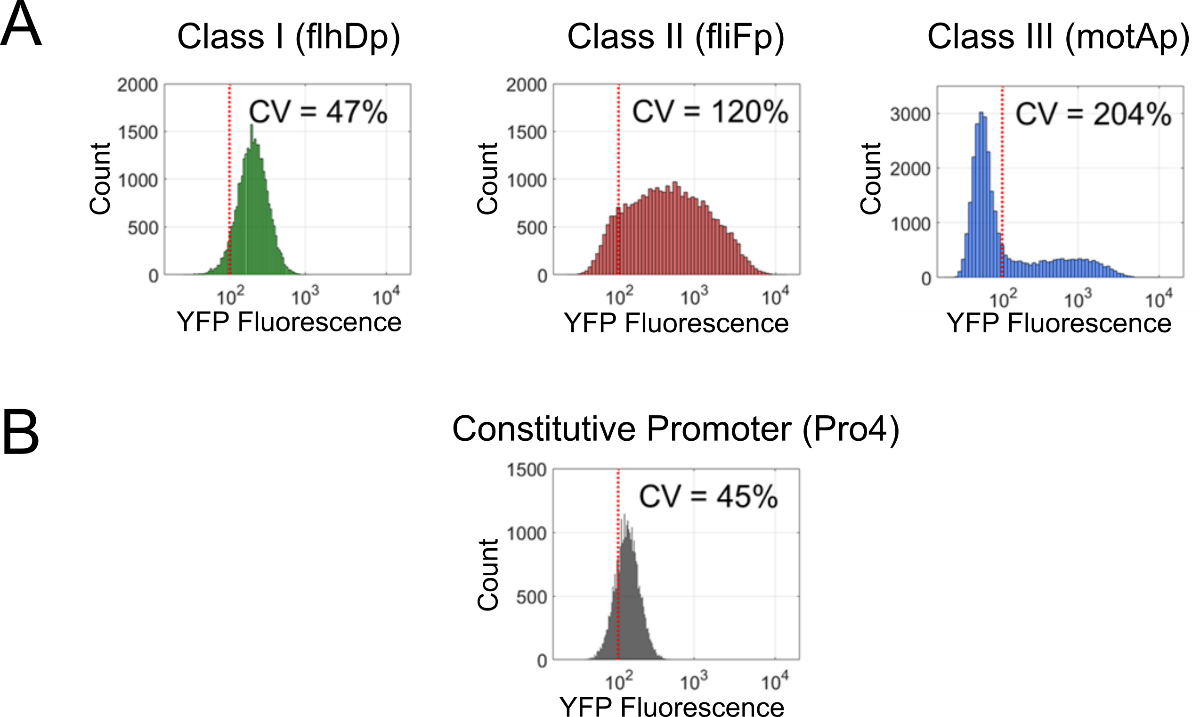

**Fig. S3. Flow cytometry analysis of flagellar gene expression for cells grown in liquid culture.** (A) Typical fluorescence distribution of Class I, II and III promoter reporters. Shown is the fluorescence distribution of cells harboring the flhD, fliF or motA promoters fused to YFP. Cells were grown in liquid culture and measured via flow cytometry as described above. Coefficient of variation (CV) values are for each fluorescence distribution. Dashed red line indicates threshold for cellular auto-fluorescence defined as the mean+2×standard deviation of flow cytometry signal from cells without any fluorescent reporters. (B) The fluorescence distribution of a constitutive promoter (Pro4) with similar mean expression to the wild-type flhD promoter. CV and dashed lines are as defined in (A).

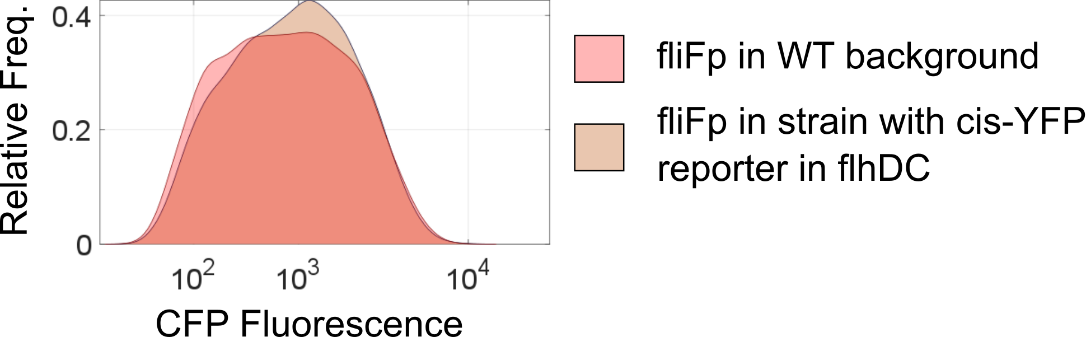

**Fig. S4. The cis-YFP reporter in the FlhDC operon does not affect Class II gene expression.** CFP fluorescence distribution for strains with a Class II (fliF)-CFP reporter with (brown) and without (red) the cis-YFP reporter in FlhDC. Cells were grown in liquid culture and measured via flow cytometry as described in “Growth Conditions”. Shown is a kernel density estimate of the fluorescence distribution obtained using the ksdensity function in MATLAB.

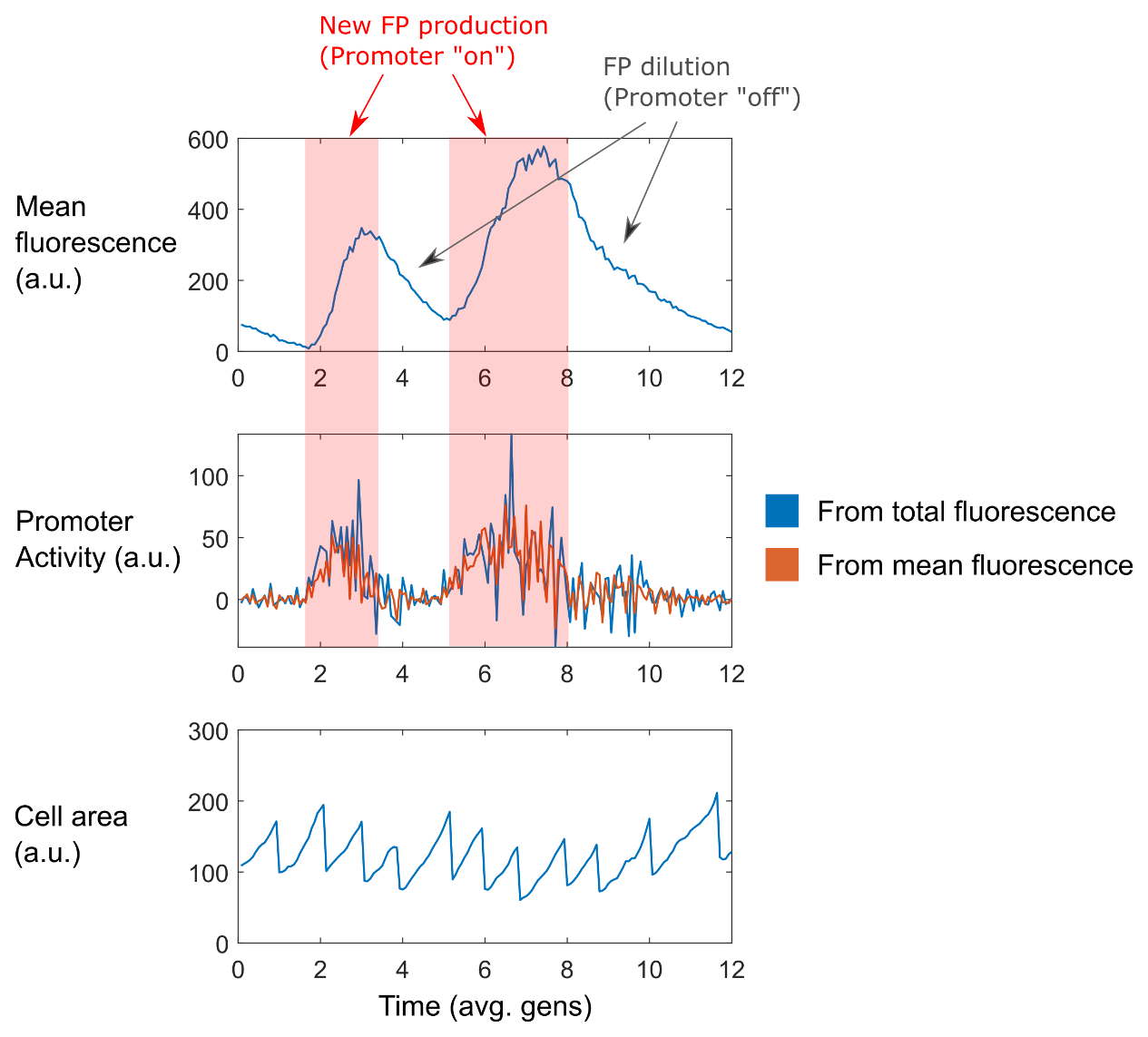

**Fig. S5. Estimation of promoter activity from time-lapse data.** Example of cell lineage trace from strain with fliFp-CFP reporter. (Top) Mean CFP fluorescence as a function of time. (Middle) Promoter activity estimated using either the total cellular fluorescence (i.e. ${P_{abs}\left( t \right)= F}_{total}\left( t \right)-F_{total}\left( t-1 \right)$, blue trace) or mean cellular fluorescence and growth rate ($P_{rel}\left( t \right)= \frac{1}{A}\frac{dF_{total}}{dt}=C\frac{1}{A}\frac{dA}{dt}+\frac{dC}{dt}$, orange trace) as described above in “Estimation of promoter activity”. Note that the two methods give different values since $P_{abs}\left( t \right)={\frac{1}{A}\times P}_{rel}\left( t \right)$ but the flagellar gene pulses are largely robust to this difference. Here we show the promoter activity estimated without applying any smoothing on the original fluorescence time-lapse data. In our analyses, we typically apply a mild Savitzky-Golay filter on the raw fluorescence trace to reduce the noise that arises when taking the numerical derivative. (Bottom) Cell area as a function of time. Red shaded region indicates time points where new FPs are created—i.e. when the promoter is “on”.

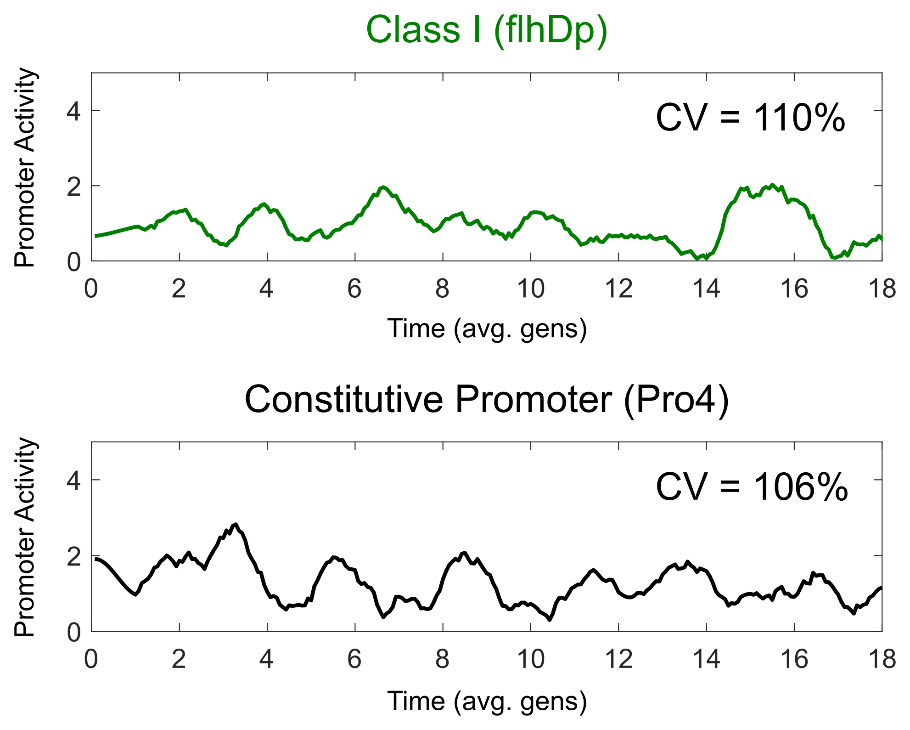

**Fig. S6. Class I (flhDp) promoter activity resembles a constitutive promoter.** Example of cell lineage trace from strain with flhDp-YFP reporter (top, green) or a constitutive promoter (Pro4-YFP) (bottom, black). The promoter activity traces were normalized by the mean promoter activity of each trace to highlight the variance relative to the mean. The CV value indicates the coefficient of variation for each strain obtained from multiple (n=20) traces.

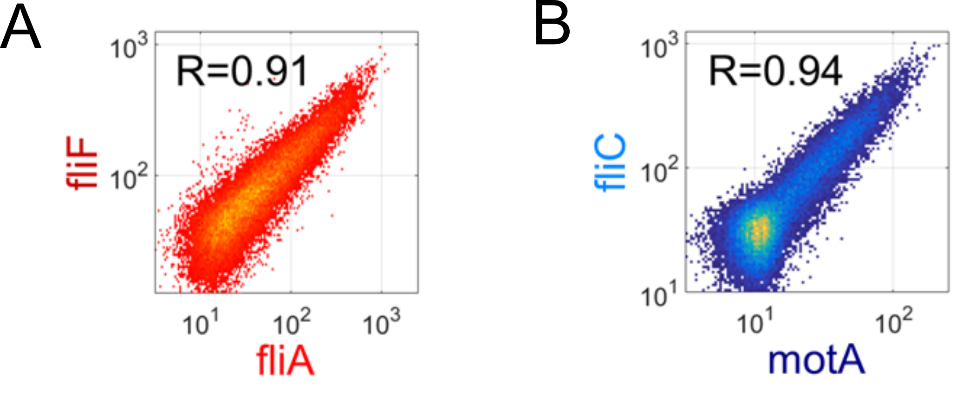

**Fig. S7****. Promoters within the same class show high correlation.** Typical fluorescence distribution in cells harboring two Class II promoter reporters (red) or two Class III promoters (blue), measured via flow cytometry. The R value indicates the Pearson correlation coefficient. (A) A 2D density plot of YFP and CFP fluorescence in cells harboring both fliAp-YFP and fliFp-CFP reporters. (B) A 2D density plot of YFP and CFP fluorescence in cells harboring both motAp-YFP and fliCp-CFP reporters.

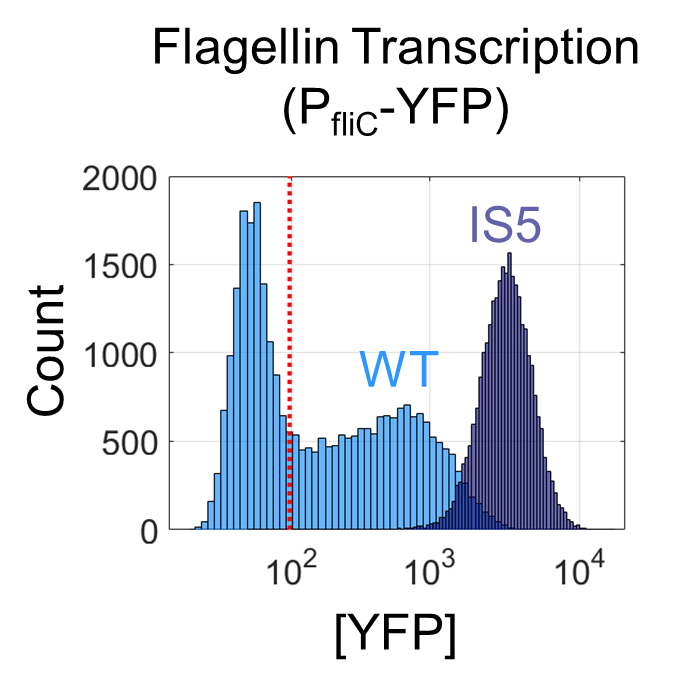

**Fig. S8****. An insertion element mutation in the FlhDC regulatory region results in high homogenous expression of flagellar genes.** Distribution of fluorescence in strains harboring PfliC-YFP reporter with (dark blue) and without (light blue) the insertion element mutation (“IS5”). Fluorescence was measured via flow cytometry. Wild type strains (WT, light blue) show large heterogeneity while insertion element mutants (IS5, dark blue, see SI text) express flagellin homogeneously. Dashed red line indicates the mean+2σ threshold for cellular auto-fluorescence (as previously described).

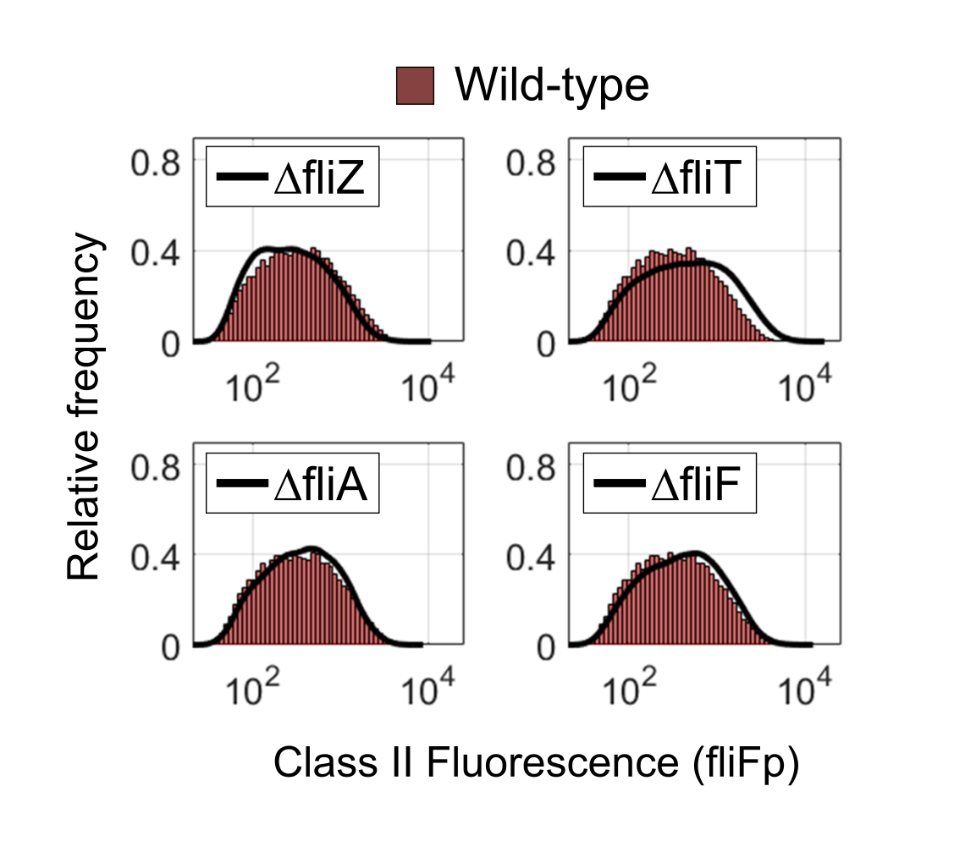

**Fig. S9****. Effect of flagellar gene deletions on Class II pulses.** Class II reporter (fliFp) fluorescence distribution for WT (red, bar histogram) and mutants (black, line). Cells were grown in liquid culture as described above in “GROWTH CONDITIONS”, and fluorescence was measured via flow cytometry. In-frame knockouts of the flagellar genes were generated using the techniques described in “Scarless Chromosomal Engineering”.
